# Adenosine impairs T cell function via activation of purine salvage pathway and AMP-induced inhibition of pyrimidine nucleotide biosynthesis

**DOI:** 10.64898/2026.09.07.749884

**Authors:** Karolina Losenkova, Ursula J. Von Danwitz, Nora Kreisig, Samuel Svärd, Julia S. Baier, Corinna Weiler, Anton Zavialov, Neil J. Thomson, Christa E. Müller, Osman Gani, Sirpa Jalkanen, Akira Takeda, Gennady G. Yegutkin

## Abstract

Extracellular adenosine (ADO) is a well-established immune checkpoint mediator suppressing T cells via activation of A_2A_ receptors, yet intracellular mechanisms of ADO action remain less understood. Here, we show that activated human T cells transport extracellular ADO through equilibrative nucleoside transporter 1 (ENT1) and metabolize it through ATP salvage pathway. The generated ADO metabolite AMP inhibits UMP synthase (UMPS), the rate-limiting enzyme of *de novo* pyrimidine biosynthesis, leading to diminished ATP production via conventional bioenergetic pathways, increased apoptosis, and impaired effector functions of T cells. Flow cytometric and single-cell transcriptomic analyses further revealed that ADO restrains the exit of naïve T cells from quiescence. All effects of ADO were phenocopied by treating T cells with the inhibitors of UMPS and reversed after pharmacological blockage of ENT1 or uridine supplementation. Additional molecular docking, molecular dynamics simulations, and free-energy analyses indicate that AMP can occupy the orotidine 5’-monophosphate (OMP) decarboxylase catalytic site of UMPS with the same anchoring contacts as substrate OMP and product UMP, providing a structural rationale for direct AMP-mediated inhibition of pyrimidine biosynthesis. These findings demonstrate that immunosuppressive effects of ADO extend beyond canonical adenosinergic signaling and involve cellular ADO uptake and modulation of purine and pyrimidine metabolism.

## INTRODUCTION

ATP and other purines are present in virtually all cells and tissues, where they exert biological effects through intracellular and extracellular mechanisms. Intracellular ATP plays a crucial role for the maintenance of cell proliferation and other energy-requiring processes ^1, 2^, whereas extracellular ATP acts as a signalling molecule mediating inflammatory, vasoactive, and angiogenic effects through a series of G protein-coupled P2Y and ligand-gated P2X receptors ^3, 4^. In turn, ATP metabolite adenosine (ADO) plays a counteracting role in mitigating inflammation through the activation of G protein-coupled adenosine receptors (AR) ^3, 5^. Based on these premises, multiple preclinical and clinical studies are underway aiming to target metabolic and signaling pathways of ADO in the tumor microenvironment (TME) by using antibodies and small-molecule inhibitors blocking the activities of key nucleotide-inactivating ectoenzymes CD39 and CD73, in combination with A_2A_R antagonists and other immune checkpoint inhibitors ^4, 6, 7^.

Along with canonical AR-dependent pathways, ADO can exert multiple biological effects via intrinsic receptor-independent mechanisms ^6, 7, 8^. Specifically, cellular uptake of ADO via equilibrative nucleoside transporter 1 (ENT1) and intracellular metabolism was shown to contribute to the control of vascular permeability, inflammation and angiogenesis ^9, 10^, maintenance of erythropoiesis and myelopoiesis ^11^, pain relief ^12^, inhibition of platelet aggregation ^13^, and invasion and proliferation of cancer cells ^14, 15, 16^. Recent studies also demonstrated the ability of ADO and/or its phosphorylated precursors ATP and AMP to inhibit T-cell receptor (TCR) activation and trigger T cell apoptosis ^8, 17, 18, 19, 20^. Blockage of the ATP-ADO axis using anti-CD39 nanobody ^18^, CD39- and CD73-inhibiting antisense oligonucleotides ^17^, or inhibitors of ENT1 and adenosine kinase (ADK) ^8, 17, 20^ restored cytokine production and proliferation of CD8^+^ T cells ^8, 17, 18, 20^, and rescued T cells from ATP-induced apoptosis ^17^. The latter observations are particularly relevant in view of the central role of T cells in the control of adaptive immune responses against cancer and other pathogens ^21, 22^. In addition to producing ATP via conventional glycolytic and oxidative phosphorylation (OxPhos) pathways, rapidly proliferating T cells are relying on continuous synthesis of ATP and other purine and pyrimidine nucleotides through *de novo* and salvage pathways to enable DNA replication, RNA production, clonal expansion, and epigenetic remodeling (Fig. 1a) ^23, 24, 25^. Furthermore, in nutrient-limited conditions T cells can utilize uridine ^25^, as well as inosine ^24^ as substrates for ATP production and gluconeogenesis via their conversion into ribose-1-phosphate to fulfil energy requirements.

**Fig. 1.**
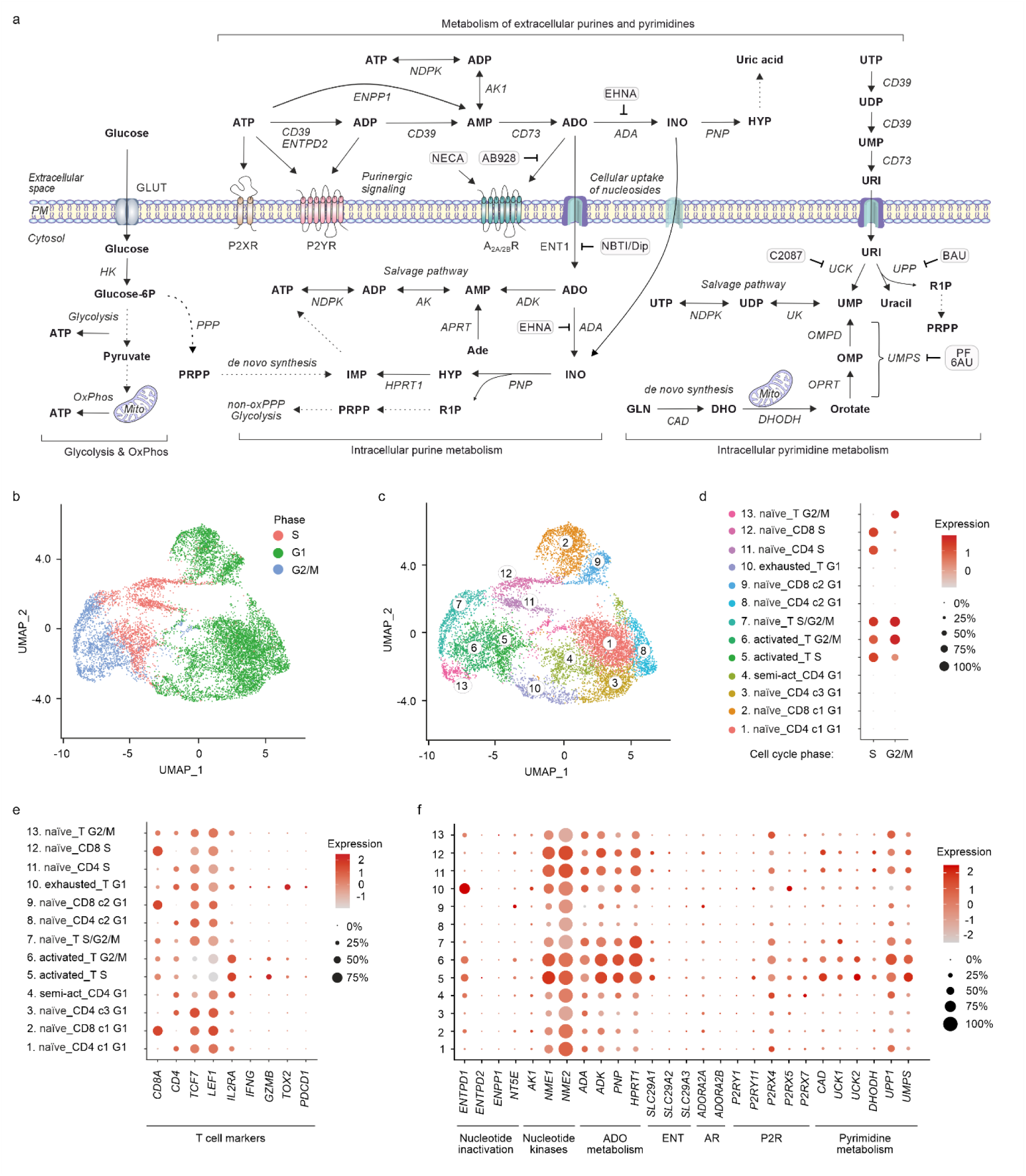
Compartmentalization of ADO biochemistry and scRNA-seq analysis of major components of purine and pyrimidine metabolism in human T cells activated by dendritic cells. (a) A schematic view of cellular bioenergetics and nucleotide metabolism. Along with canonical ATP biosynthesis from glucose via glycolysis and oxidative phosphorylation (*OxPhos*), other pathways contribute to energy homeostasis, including synthesis of ATP and other nucleotides through the *de novo* and salvage pathways. Cellular purine turnover includes: (i) the release of ATP and activation of P2X and P2Y receptors; (ii) metabolism of extracellular purines; (iii) activation of ADO receptors (AR, encoded by *ADORA* gene); and (iv) cellular ADO uptake via equilibrative nucleoside transporter 1 (ENT1, *SLC29A1*). General schemes of extracellular purine metabolism have included a role for the ecto-nucleoside triphosphate diphosphohydrolase-1 (NTPDase1, known as CD39, gene name *ENTPD1*), NTPDase2 (*ENTPD2*), ecto-nucleotide pyrophosphatase/phosphodiesterase-1 (ENPP1), ecto-5’-nucleotidase (CD73, *NT5E*), adenosine deaminase (ADA), purine nucleoside phosphorylase (PNP), as well as ATP-regenerating ectoenzymes of the nucleotide diphosphate kinase (NDPK, *NME*) family and adenylate kinase-1 (AK1). Following cellular uptake, ADO can be inactivated via inosine (INO) to hypoxanthine (HYP), or phosphorylated to AMP and further to ADP and ATP. Other nucleosides are generated from the corresponding extracellular nucleotides and taken up by the cells. For sake of clarity, only metabolism of uridine (URI) via *de novo* pyrimidine biosynthesis and salvage pathway is shown. Sites of manipulation of metabolic pathways by using UMP synthase (UMPS) inhibitors pyrazofurin (PF) and 6-azauridine (6AU), ENT1 inhibitors NBTI and dipyridamole (NBTI/Dip), ADA inhibitor EHNA, uridine-cytidine kinase (UCK) inhibitor Compound 20874830-2 (*C2087*), and uridine phosphorylase-1 (UPP1) inhibitor benzylacyclouridine (BAU) are indicated by blunt-ended arrows. Dotted lines indicate indirect links between two metabolites. Other abbreviations: Ade, adenine; ADK, adenosine kinase; APRT, adenine phosphoribosyltransferase; CAD, carbamoyl-phosphate synthetase; DHODH, dihydroorotate dehydrogenase; DHO, dihydroorotate; GLN, glutamine; GLUT, glucose transporter; HK, hexokinase; HPRT1, hypoxanthine phosphoribosyltransferase 1; Mito, mitochondria; non-oxPPP, non-oxidative pentose phosphate pathway; OMP, orotidine 5’-monophosphate; OMPD, OMP decarboxylase; OPRT, orotate phosphoribosyl transferase; PM, plasma membrane; PRPP, Phosphoribosyl pyrophosphate; R1P, ribose-1-phosphate; UK, uridylate kinase. (b-e) Human T cells were activated by co-culturing for six days with the antigen-presenting CD1c^+^ dendritic cells (DC) isolated from human lymph nodes. Two independent datasets were integrated. A uniform manifold approximation and projection (UMAP) plots identified 13 distinct clusters, which were numbered according to the T cell cycle phase (S/G1/G2/M) (b) or the expression of CD4, CD8, and other T cell markers (c). Dot plots show differential gene expression profiles of the naïve, exhausted, semi-activated (semi-act), and activated T cells. The results are expressed on a pseudocolor scale (log2(FC)), with the size of the dot representing the percentage of T cells maintained in a certain phase of cell cycle (d) or expressing the indicated T cell marker (e). (f) The expression of genes encoding key metabolic enzymes, as well as ENTs and nucleotide- and ADO-specific receptors.

Here we show that, upon activation, human T cells reprogram metabolic machinery via down-regulation of ecto-5’-nucleotidase/CD73 and up-regulation of CD39, adenosine deaminase (ADA) and other purine-converting enzymes, as well as ENT1. These shifts allow activated T cells to avoid accumulation of ADO in their vicinity and simultaneously utilize it as a fuel for replenishing intracellular ATP stores. Moreover, we found that phosphorylation of cell-incorporated ADO via AMP into ADP and ATP, and concurrent AMP-mediated inhibition of the rate-limiting enzyme of pyrimidine nucleotide biosynthesis UMP synthase (UMPS) constitutes a major mechanism that suppresses energy metabolism, effector functions and viability of activated T cells, and also hinders the exit of naïve T cells from quiescence. These data provide a novel insight into the intrinsic mechanisms of ADO regulating T cell function, viability, and nucleotide homeostasis.

## RESULTS

### 1. Transcriptomic analysis reveals dynamic shifts in metabolic and signaling pathways of purines and pyrimidines in human T cells upon activation

In the first part of this study, we assessed activation-induced shifts in T cell metabolism, with particular emphasis on key enzymes and transporters involved in purine and pyrimidine metabolism (Fig. 1a). T cells were isolated from blood of healthy donors, activated by co-culturing for six days with allogenic CD1c^+^ dendritic cells (DC) isolated from the human lymph nodes, and subjected to single-cell RNA sequencing (scRNA-seq) analysis. Unbiased clustering revealed distinct clusters, distinguished by different cell cycle phases (S/G1/G2/M), and by the expression of CD4, CD8, and markers of naïve (*TCF7*, *LEF1*), activated (*IL2RA*, *IFNG*, *GZMB*), and exhausted (*TOX2*) T cells (Fig 1b-e). Additional analysis revealed that all activated T cells in either S-phase (cluster 5) or G2/M-phase (cluster 6) displayed high levels of enzymes involved in the metabolism of extracellular and/or intracellular nucleotides (CD39/*ENTPD1,* NDPK-A/*NME1,* NDPK-B/*NME2*), and purine (*ADA, ADK, PNP, HPRT1*) and pyrimidine (*CAD, UCK1/2, UPP1, UMPS*) nucleosides, as well as ENT1 (*SLC29A1*), as compared with naïve and semi-activated T cells. Cluster 10 exhibited the signature of exhausted T cells, characterized by high expression of PD-1 (*PDCD1*), *TOX2*, and *ENTPD1* (Fig. 1e). T cells also express *NT5E*, with relatively high expression of this ecto-nucleotidase detected in the naïve CD8^+^ T cells (clusters 2, 9, 12).

We also utilized a public scRNA-seq dataset of human T cells incubated with T-Activator CD3/CD28 Dynabeads ^26^. Unbiased clustering identified 9 clusters, distinguished based on the expression of CD4, CD8, and other T cell markers (Supplementary Fig. 1a). As in the case of DC-primed T cells, CD3/CD28-stimulated CD8^+^ and CD4^+^ T cells express high levels of genes encoding NDPK, ADK, PNP, HPRT1, UMPS, CAD, UCK1/2, and ENT1 (Supplementary Fig. 1b). In this dataset, high CD39/*ENTPD1* was detected only in CD4^+^/FOXP3^+^ T regulatory (T_reg_) cells, but not in activated T cells (Supplementary Fig. 1b), which could be due to relatively short 16-hour stimulation, insufficient for up-regulation of CD39 on activated T cells ^19, 27^. While a detailed characterization of signalling pathways lies beyond the scope of this study, we analyzed the expression of receptors mediating biological effects of nucleotides and ADO. Both DC-primed (Fig. 1f), and CD3/CD28-stimulated (Supplementary Fig. 1b) T cells showed similar dynamics of up-regulation of nucleotide-selective receptors of P2Y_11_ (*P2RY11*), P2X4 (*P2RX4*), P2X5 (*P2RX5*), and P2X7 (*P2RX7*) subtypes. Naïve CD8^+^ T cells and, to a lesser extent, naïve CD4^+^ T cells express A_2A_R (*ADORA2A*) (Fig. 1f, Supplementary Fig. 1b), whereas other AR subtypes (*ADORA2B*, *ADORA1*, and *ADORA3*) were expressed at low or undetectable levels in all T cell subsets.

Overall, despite some variations among two models, scRNA-seq analysis of activated T cells revealed similar pattern of changes in the expression profiles of genes encoding key components of purine and pyrimidine turnover. Upon activation, T cells down-regulate A_2A_R (*ADORA2A*) and CD73 (*NT5E*), while up-regulating other enzymes that control extra- and intracellular levels of nucleotides, as well as ENT1 (*SLC29A1*). Based on these data, and also taking into account previous data on dynamic changes in CD39 and CD73 expression on T cells incubated with anti-CD3/CD28 beads from 0 to 7 days ^19, 27^, in the following experiments we extended the period of T cell stimulation with the beads up to 96 hours and used this experimental model for assessing the role of ADO in T cell function.

### 2. Cellular purine turnover is augmented during T cell activation through concerted up-regulation of key purine-converting enzymes and ENT1-mediated ADO uptake

Next, we sought to characterize activation-induced shifts in catalytic activities and expression levels of key enzymes of purine metabolism. T cells were incubated without and with anti-CD3/CD28 beads and defined in all subsequent experiments as “resting” and “activated” T cells, respectively. The activities of ecto-nucleotide hydrolases and kinases were assayed by incubating T cells with ^3^H-labeled ADP and AMP in the absence or presence of γ-phosphate-donating ATP. Tracer nucleotide substrates and their ^3^H-metabolites were separated by thin layer chromatography (TLC), and quantified by scintillation β-counting (Fig. 2a) or developed by autoradiography (Fig. 2b). As shown in Fig. 2b, when resting (lane 1) and activated (lane 2) T cells were incubated with [^3^H]ADP as a preferred substrate for CD39, a clearly detectable band corresponding to [^3^H]AMP was revealed only in the latter case. In the presence of ATP, [^3^H]ADP was converted into [^3^H]ATP, with high NDPK activity detected in activated (lane 4) and to a lesser extent, in resting (lane 3) T cells. Likewise, joint addition of [^3^H]AMP and ATP to the activated (lane 10), but not resting (lane 9), T cells caused their transphosphorylation into ^3^H-labeled ADP/ATP through sequential AK and NDPK reactions. Quantitative analysis confirmed significantly increased CD39, NDPK and AK activities in activated T cells (Fig. 2a). Conversely, the rate of CD73-mediated hydrolysis of [^3^H]AMP to [^3^H]ADO was decreased during T cell activation (Fig. 2a, and Fig. 2b, lanes 7-8).

**Fig. 2.**
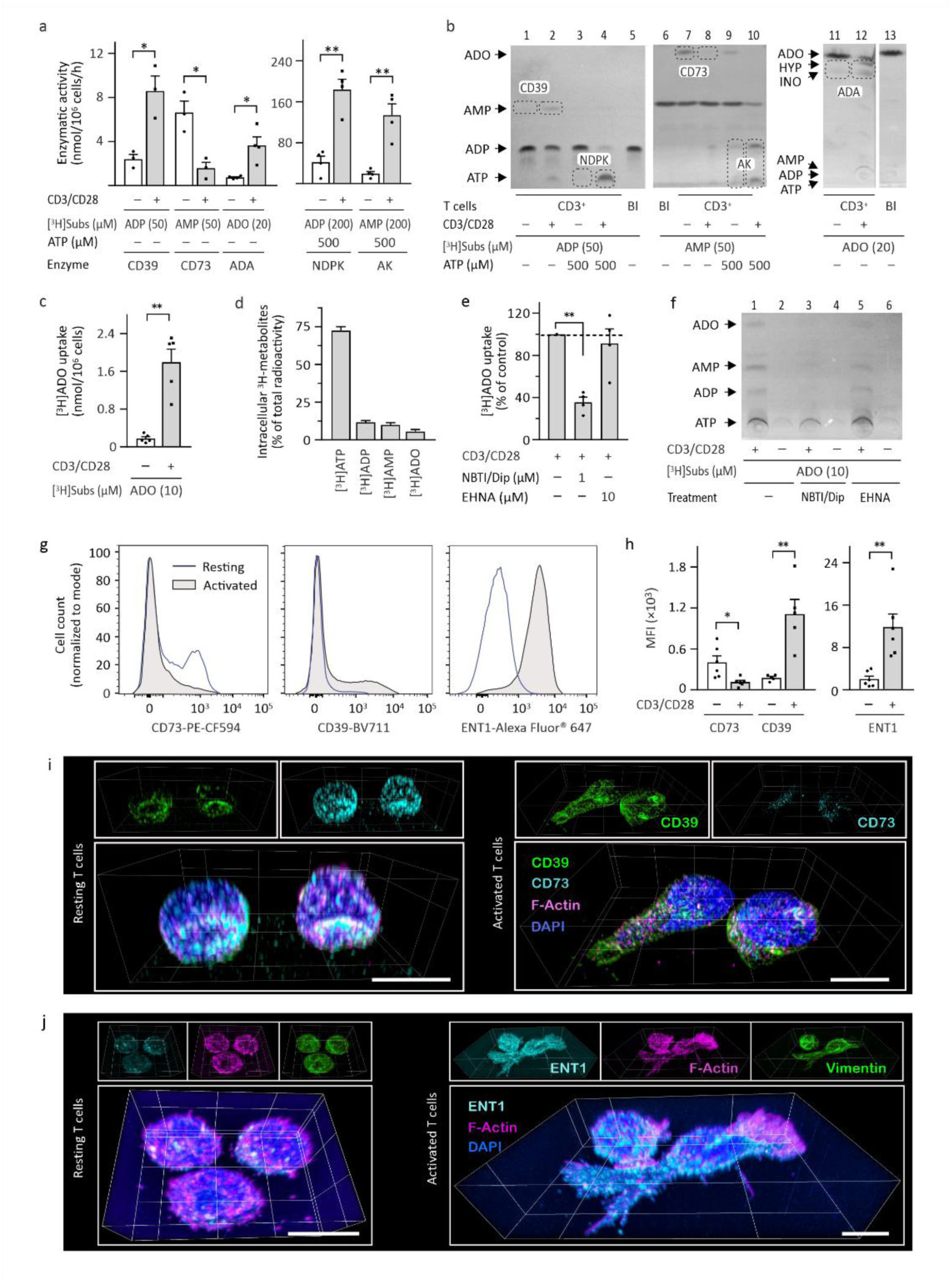
Activated T cells reprogram purine metabolism through down-regulation of CD73 and up-regulation of CD39, ADA, other purinergic enzymes, as well as ENT1. T cells were incubated for 96 hours without or with anti-CD3/CD28 beads, and subsequently incubated with [^3^H]ADP, [^3^H]ADP plus ATP, [^3^H]AMP, [^3^H]AMP plus ATP, and [^3^H]ADO as respective substrates for CD39, NDPK, CD73, AK, and ADA. ^3^H-labeled substrates and their metabolites were separated by TLC using two different solvent systems employed for the measurement of nucleotide-converting and ADA activities. (a) Specific enzymatic activities were expressed as nanomoles of ^3^H-substrates inactivated (in the case of CD39, CD73 and ADA) or phosphorylated (for NDPK and AK) by one million cells per one hour. (b) Representative autoradiographic images of the patterns of [^3^H]ADP, [^3^H]AMP, and [^3^H]ADO metabolism by T cells. The blanks (Bl) show the radiochemical purities of tracer substrates. Arrows indicate the position of nucleotide and nucleoside standards. Resting and activated T cells were incubated for 60 min with 10 μM [^3^H]ADO, washed, lysed, separated by TLC, and analyzed by scintillation β-counting for the amounts of [^3^H]ADO and its intracellular ^3^H-metabolites. The results are expressed as nanomoles of [^3^H]ADO taken up by one million cells (c), or as a relative amount of ^3^H-labeled ADO, AMP, ADP, and ATP (d). T cells were pre-treated with a mixture of ENT1 inhibitors, NBTI and dipyridamole (NBTI/Dip), or ADA inhibitor EHNA, prior to addition on [^3^H]ADO. (e) The results are expressed as a percentage of radioactivity determined in control T cells. (f) Representative autoradiographic image of the pattern of intracellular [^3^H]ADO metabolism by resting and activated T cells. The expression levels of ecto-nucleotidases and ENT1 were assayed by flow cytometry. (g) Representative histograms are shown for the resting (*Rest*) and activated (*Act*) T cells stained with anti-CD73, anti-CD39, and anti-ENT1 antibodies. (h) Mean fluorescence intensities (MFI) of CD39^+^, CD73^+^ and ENT1^+^ T cells were determined by FlowJo. The results are presented as scatter plots with bars (mean + SEM). *P<0.05 and **P<0.01, determined by Student’s t test (paired, two-tailed). Resting and activated T cells were co-stained with anti-CD73 and anti-CD39 (i), or anti-ENT1 and anti-vimentin (j) antibodies, together with the F-actin binding drug Alexa Fluor® 546-phalloidin. Z-stacks were captured using a confocal microscope and presented as 3D images. Lower images show the merged images with nuclei counterstained with DAPI. Scale bars, 6 µm.

For the measurement of ADA activity, we used [^3^H]ADO as an initial substrate and a particular solvent system that enabled better separation of [^3^H]ADO and its ^3^H-metabolites ^28^. Addition of [^3^H]ADO to activated T cells caused its deamination into [^3^H]inosine and further to [^3^H]hypoxanthine through sequential ADA and PNP reactions, whereas resting T cells displayed relatively low ADA activity (Fig. 2a, and Fig. 2b, lanes 11-12). Additional immunomagnetic separation of CD3^+^ T cells into CD4^+^ and CD8^+^ T cells revealed a similar pattern of opposite down-regulation of CD73 and up-regulation of ADA activities on both T cell subtypes upon their activation (Supplementary Fig. 2).

To elucidate a link between extra- and intracellular ADO metabolism, T cells were incubated with 10 μM [^3^H]ADO and assayed by TLC for the amount of cell-incorporated [^3^H]ADO and its ^3^H-metabolites. The ability of activated T cells to take up [^3^H]ADO was ∼7-10-fold higher compared to the resting controls (Fig. 2c). Additional analysis has shown that, following its transport, [^3^H]ADO was phosphorylated into [^3^H]ATP, which accounted for >70% of the total radioactivity (Fig. 2d). For blocking [^3^H]ADO uptake, we used a potent inhibitor of ENT1, NBTI ^29^, together with dipyridamole, an antithrombotic drug that mediates pleiotropic biological effects, including inhibition of ENT1 ^13, 30^. Treatment of activated T cells with NBTI and dipyridamole prior to addition of [^3^H]ADO significantly diminished the amount of cell-incorporated radioactivity, whereas ADA inhibitor EHNA did not affect intracellular [^3^H]ADO metabolism (Fig. 2e,f). Consistent with radio-TLC data, flow cytometric analysis confirmed increased cell surface expression of CD39 and ENT1 and decreased CD73 expression on activated T cells (Fig. 2g,h).

Additional three-dimensional (3D) confocal microscopic imaging demonstrated punctate distribution of both CD73 and CD39 across the plasma membrane of resting T cells, while activated T cells were characterized by increased CD39 immunoreactivity and relatively faint CD73-specific fluorescence (Fig. 2i). Cellular shape and ENT1 distribution were also visualized by co-staining T cells with the anti-ENT1 antibody, together with a marker of intermediate filaments vimentin, and the F-actin-binding drug Phalloidin. Resting T cells displayed well-structured cytoskeletal and filamentous networks organized into thick cortical rings along the cell periphery with preferential accumulation of F-actin fibers in fine plasma membrane protrusions (microvilli). Upon activation, T cells undergo structural rearrangements, manifested in the polarization of cell body from spheroid to flattened shape, pseudopod protrusion, F-actin polymerization, stress fiber formation at the leading edge, as well as increased cell surface expression of ENT1 (Fig. 2j, right). These confocal microscopy data were corroborated by flow cytometric analysis showing markedly increased size and granularity of activated T cell (Supplementary Fig. 3).

Collectively, by using a radio-TLC approach, in combination with flow cytometric and immunofluorescent analyses, we characterized the whole pattern of cellular ADO turnover in human T cells, which includes: (a) metabolism of extracellular nucleotides and nucleosides via a network of purine-inactivating (CD39, CD73, ADA, and PNP) and ATP-regenerating (NDPK and AK) ectoenzymes, (b) cellular influx of ADO via ENT1, and finally (c) conversion of ADO into intracellular AMP and further to ADP and ATP through sequential phosphotransfer reactions. Except for diminished expression of CD73, other components of purine turnover cascade significantly up-regulated in T cells upon activation.

### 3. ADO inhibits T cell bioenergetics via cellular uptake through ENT1 and rewiring ATP production from glycolysis and OxPhos to purine salvage pathway

Considering the above data on the ability of T cells to uptake [^3^H]ADO and phosphorylate it into [^3^H]ATP, we assessed the effect of ADO on cellular ATP levels and energy metabolism. T cells were incubated with anti-CD3/CD28 beads, ADO and other drugs, as outlined in Fig. 3a. Bioluminescent analysis revealed a two-fold increase in intracellular ATP during T cell stimulation, equal to 0.99 ± 0.10 and 1.96 ± 0.16 nanomoles of ATP per million of resting and activated T cells, respectively (Supplementary Fig. 4a). When ATP concentrations were expressed as nanomoles per milligram of cellular protein, rather moderate, albeit statistically significant, 20% increase in ATP was detected in activated T cells (Supplementary Fig. 4b). These results, together with data on the increased complexity of activated T cells (Fig. 2j, Supplementary Fig. 3), suggest that T cells maintain ATP at certain characteristic levels, which rise upon T cell activation proportionally to their size, granularity and total protein content.

**Fig. 3.**
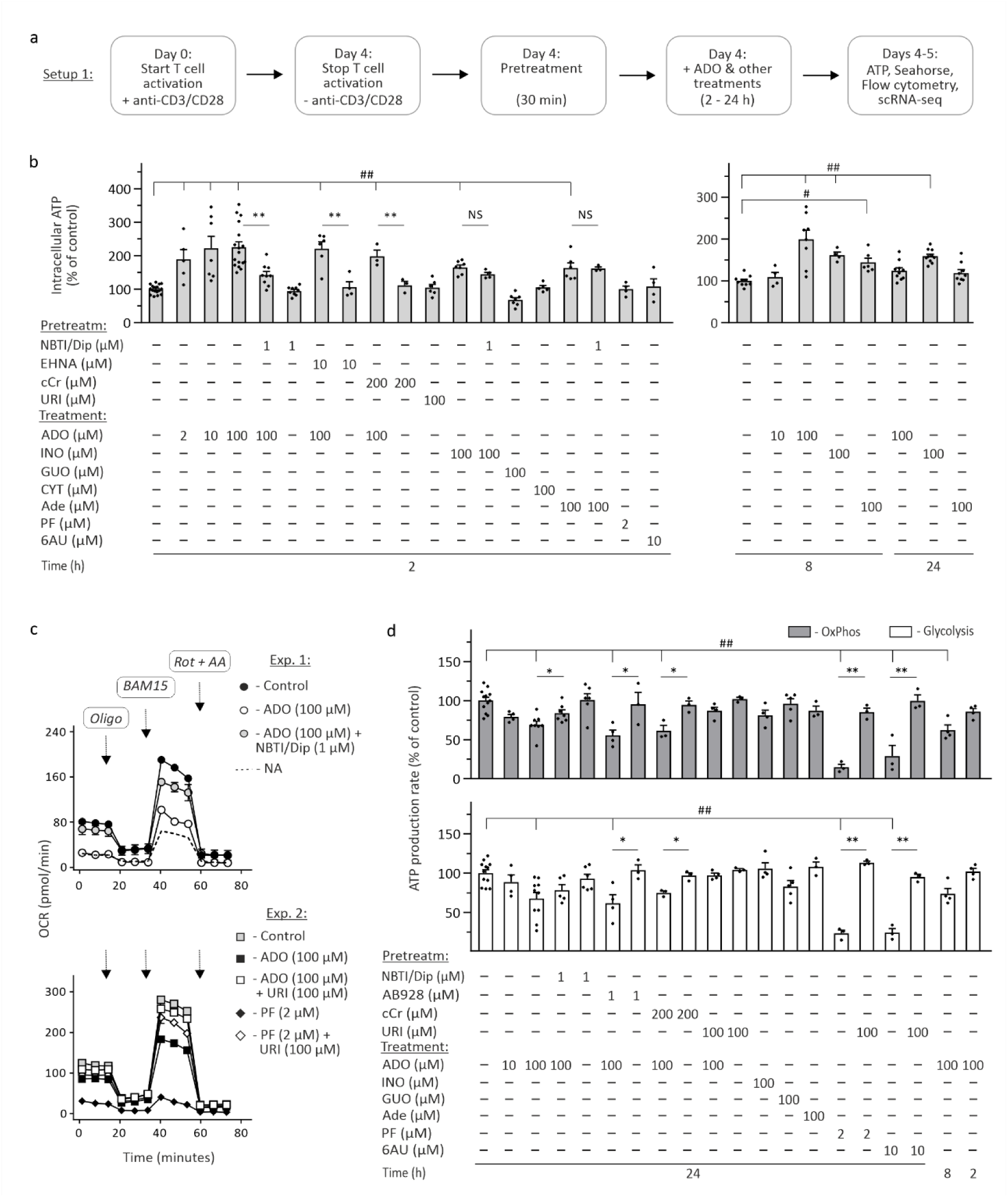
ADO modulates energy metabolism and ATP production by activated T cells via cellular uptake through ENT1 and inhibition of pyrimidine synthesis. (a) Workflow of the study design. T cells were stimulated for four days with anti-CD3/CD28 beads, pretreated with different inhibitors, and incubated for additional 2-24 hours with ADO and other related drugs. (b) Bioluminescent analysis of global intracellular ATP levels in activated T cells subjected to different treatments. The results are plotted as the percentage of ATP concentrations measured in control vehicle-treated cells. T cell bioenergetics was determined using the Agilent Seahorse XF Cell Mito Stress Kit. Measurements of oxygen consumption rates (OCR) after sequential injections of oligomycin A (*Oligo*), the mitochondrial uncoupler BAM15, and mitochondrial inhibitors rotenone and antimycin A (*Rot + AA*) enable the calculation of ATP production via mitochondrial OxPhos. (c) Representative time courses of oxygen consumption by control and treated activated T cells and also by non-activated (*NA*) T cells from two independent experiments are shown. (d) Simultaneous measurement of the rates of oxygen consumption and extracellular acidification allowed to determine cellular ATP production through OxPhos (upper plot) and glycolysis (lower plot). The results are expressed as a percentage of control (mean + SEM). Statistical significances in panels b and d were determined by one-way ANOVA with Dunnett’s multiple comparison test (^#^P<0.05 and ^##^P<0.01) or unpaired Student’s t-test (*P<0.05 and **P<0.01).

Incubation of activated T cells with the increasing concentrations of ADO (2–100 μM) temporarily increased intracellular ATP, which reached a maximum ∼2–3-fold peak after two hours. This ATP spike was prevented by pretreating T cells with ENT1 inhibitors NBTI and dipyridamole, whereas ADA inhibitor EHNA did not reverse ADO-induced rise in cellular ATP. ATP levels were also increased by ∼50% after a 2-hour incubation of activated T cells with 100 μM inosine or purine nucleobase adenine, but these effects were not abolished by NBTI and dipyridamole. Cellular ATP remained elevated after 24 hours of incubation of T cells with inosine, but not with ADO. Incubation of activated T cells with other purine (guanosine) and pyrimidine (uridine, cytidine) nucleosides did not affect ATP concentrations (Fig. 3b). No changes in cellular ATP were observed after incubation of resting T cells with 100 μM ADO (Supplementary Fig. 4c). To further examine the effect of ADO on T cell energy metabolism, we took advantage of the Agilent Seahorse metabolic assay. Real-time measurements of ATP production showed that the glycolysis-to-OxPhos ratio was increased during T cell stimulation (Supplementary Fig. 5a). These data are consistent with the current view that, upon antigen stimulation and TCR activation, T cells rewire metabolism from a quiescent state reliant on OxPhos to a highly glycolytic ATP-generating phenotype to meet the rising energy demands ^21, 22, 31^. Incubation of activated (Fig. 3c,d), but not resting (Supplementary Fig. 5b), T cells with 100 µM ADO for 24 hours suppressed both glycolytic and mitochondrial ATP production. These inhibitory effects of ADO were reversed by pretreating T cells with NBTI and dipyridamole, but not with the A_2A/2B_R antagonist AB928. Previous studies demonstrated that creatine may act as a “molecular battery” conserving energy in the form of ATP to power T cell activities ^32^. However, the inhibitor of creatine kinase cyclocreatine (cCr) had no effect on intracellular ATP levels (Fig. 3b) and the rate of ATP production (Fig. 3d) in activated T cells, neither when added alone nor in combination with ADO. Interestingly, ADO was able to inhibit mitochondrial ATP production after 8 hours, whereas reduction of the exposure time to 2 hours was insufficient for blocking T cell bioenergetics (Fig. 3d). The effect of ADO on energy metabolism was also studied in CD4^+^ and CD8^+^ T cell subsets. ADO inhibited both glycolysis and OxPhos in activated CD8^+^ T cells and suppressed mitochondrial respiration in CD4^+^ T cells (Supplementary Fig. 6a,b). Two UMPS inhibitors, PF and 6AU, suppressed ATP production by the activated T cells even more robustly than ADO. Strikingly, supplementation of the assay buffer with 100 μM uridine reversed inhibitory effects of PF, 6AU, as well as ADO (Fig. 3c,d). No changes in ATP production rates were detected in the presence of uridine, guanosine, inosine or adenine. These data indicate that, while activated T cells efficiently utilize ADO for replenishing intracellular ATP levels, prolonged exposure to ADO results in suppression of conventional T cell bioenergetics through intrinsic mechanisms involving cellular ADO transport and inhibition of pyrimidine nucleotide biosynthesis.

### 4. ADO induces apoptosis and impairs IFN-γ production by activated T cells via cellular uptake through ENT1 and inhibition of pyrimidine nucleotide metabolism

We then evaluated the effect of ADO on the viability (Fig. 4a) and effector function (Fig. 4b) of T cells. Incubation of activated (Fig. 4a,c), but not resting (Supplementary Fig. 7), T cells with 100 µM ADO for 24 hours resulted in a significant rise in the percentage of Annexin V^+^/7AAD^+^ apoptotic cells. These cytotoxic effects of ADO were prevented by pretreating the cells with a mixture of ENT1 inhibitors NBTI and dipyridamole, and also with uridine (Fig. 4a,c). ADO also diminished the ability of activated T cells to secrete inflammatory cytokine IFN-γ, whereas blockage of ENT1 with NBTI and dipyridamole, or addition of uridine restored the impaired IFN-γ production by ADO-treated T cells (Fig. 4b,d). Pretreatment of activated T cells with ADA inhibitor EHNA or A_2A/2B_R antagonist AB928 did not prevent cytotoxic (Fig. 4c) and immunosuppressive (Fig. 4d) effects of ADO. As in the case of ADO treatment, the inhibitors of UMPS, PF and 6AU, increased apoptosis rate and inhibited IFN-γ production by activated T cells, with all these effects being reversed by uridine (Fig. 4). Noteworthy, NBTI and dipyridamole also abolished the effects of 6AU, but not PF. These findings, together with data on decreased efficacy of uridine to relieve the inhibitory effects of PF in the presence of NBTI and dipyridamole (Fig. 4c,d), indicate that ENT1 facilitates cellular uptake of ADO, uridine, and 6AU, whereas PF penetrates T cells via a different route.

**Fig. 4.**
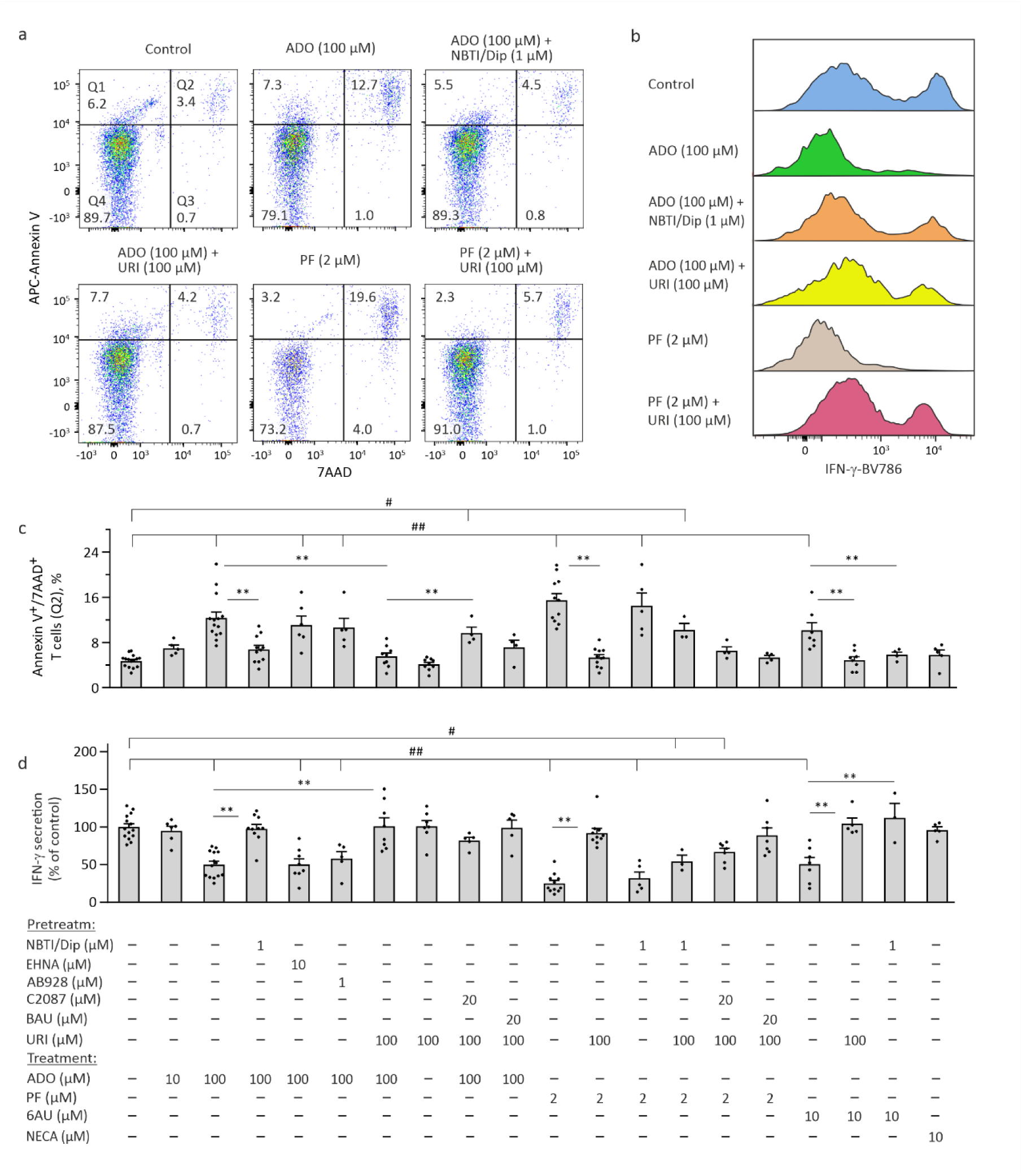
The effect of ADO and other purinergic and pyrimidinergic drugs on T cell viability and IFN-γ production by activated T cells. T cells were stimulated for four days with anti-CD3/CD28 beads, incubated for additional 24 hours with ADO and other compounds, as outlined in Figure 3a, and further assessed for cell viability and effector function by flow cytometry. (a) Representative dot plots for Annexin V versus 7AAD staining of control and treated T cells. The gating strategy and percentage of the viable cells (Q4), cells in early apoptosis (Q1), and late apoptotic or dead cells (Q2) are shown. (b) The graphs show representative histograms for the production of IFN-γ by control and treated T cells. (c,d) Quantitative analyses of the induction of apoptosis and IFN-γ production by activated T cells treated with the indicated concentrations of ADO and other drugs. The frequencies of annexin V^+^/7AAD^+^ apoptotic T cells (Q2) (c) and mean fluorescence intensities of IFN-γ^+^ T cells (d) were quantified using FlowJo (mean + SEM). Statistical significances were determined by one-way ANOVA with Dunnett’s multiple comparison test (^#^P<0.05 and ^##^P<0.01), or unpaired Student’s t-test (*P<0.05 and **P<0.01).

To further characterize the mechanism of rescuing action of uridine, activated T cells were treated with UCK inhibitor C2087 or UPP inhibitor BAU. Pretreatment of the cells with C2087, but not BAU, attenuated the ability of uridine to reverse cytotoxic effects of ADO (Fig. 4c) and suppressive effects of PF (Fig. 4d). These competitive data suggest that, upon cellular uptake via ENT1, uridine is mainly phosphorylated into UMP through UCK reaction rather than undergoes phosphorolysis to uracil by UPP (see Fig. 1a). Treatment of activated T cells with a lower dose of ADO (10 µM), other nucleosides (uridine, inosine, guanosine, or cytidine at 100 µM), the A_2A_/_2B_R agonist NECA, as well as any of the metabolic pathway inhibitors added alone did not alter T cell viability or the production of IFN–γ (Fig. 4c,d; Supplementary Fig. 8). Additional flow cytometric analysis demonstrated significantly decreased expression of Granzyme B (GranzB) in the PF-treated T cells, whereas ADO, 6AU, and other treatments did not affect the ability of activated T cells to secrete this cytotoxic molecule (Supplementary Fig. 9).

These findings indicate that ADO mediates cytotoxic and suppressive effects on activated T cells via AR-independent mechanism, which involves cellular nucleoside uptake through ENT1 and inhibition of pyrimidine nucleotide metabolism. This conclusion is supported by data showing that direct blockage of the *de novo* pyrimidine biosynthesis using UMPS inhibitors PF and 6AU also increased apoptosis and prevented IFN-γ production by activated T cells. Importantly, supplementation of the assay buffer with uridine activated alternative pyrimidine salvage pathway through UCK reaction, thus allowing activated T cells to bypass UMPS deficiency and to reverse all adverse effects of ADO, PF, and 6AU.

### 5. ADO suppresses transition of T cells from a quiescent to activated state through cellular uptake via ENT1 and inhibition of pyrimidine nucleotide biosynthesis

In a separate set of experiments, T-Activator CD3/CD28 Dynabeads, different inhibitors, ADO and other nucleosides were added successively to the freshly isolated T cells, as outlined in Fig. 5a. To assess cell proliferation, T cells were labelled with the fluorescent dye Cell Trace™ Violet (CTV) prior to addition of the beads and other treatments. Incubation of T cells with anti-CD3/CD28 beads in the presence of 100 μM ADO was accompanied by impaired proliferation of CTV-labeled T cells, with significant effects being observed even at lower (10 μM) concentration of ADO (Fig. 5b,d). Addition of 100 μM ADO to the naïve T cells also suppressed their differentiation into effector T cells, as ascertained by diminished ability to produce IFN-γ and GranzB, along with the decreased surface expression of PD-1, CD25, and ENT1 (Fig. 5c, e-i). ENT1 inhibitors (NBTI and dipyridamole) and uridine, but not A_2A_R/A_2B_R antagonist AB928, counteracted the anti-proliferative effects of ADO, restored the impaired ability of ADO-treated T cells to produce IFN-γ and GranzB, and normalized the expression levels of T cell activation markers. Similar to ADO treatment, direct blockage of *de novo* pyrimidine nucleotide biosynthesis by the inhibitors of UMPS, PF and 6AU, exerted potent immunosuppressive effects, which were rescued by adding uridine (Fig. 5d-i). Incubation of T cells with A_2A_R/A_2B_R agonist NECA, uridine, inosine (Fig. 5), as well as guanosine and cytidine (data not shown), did not affect any aspects of T cell activation.

**Fig. 5.**
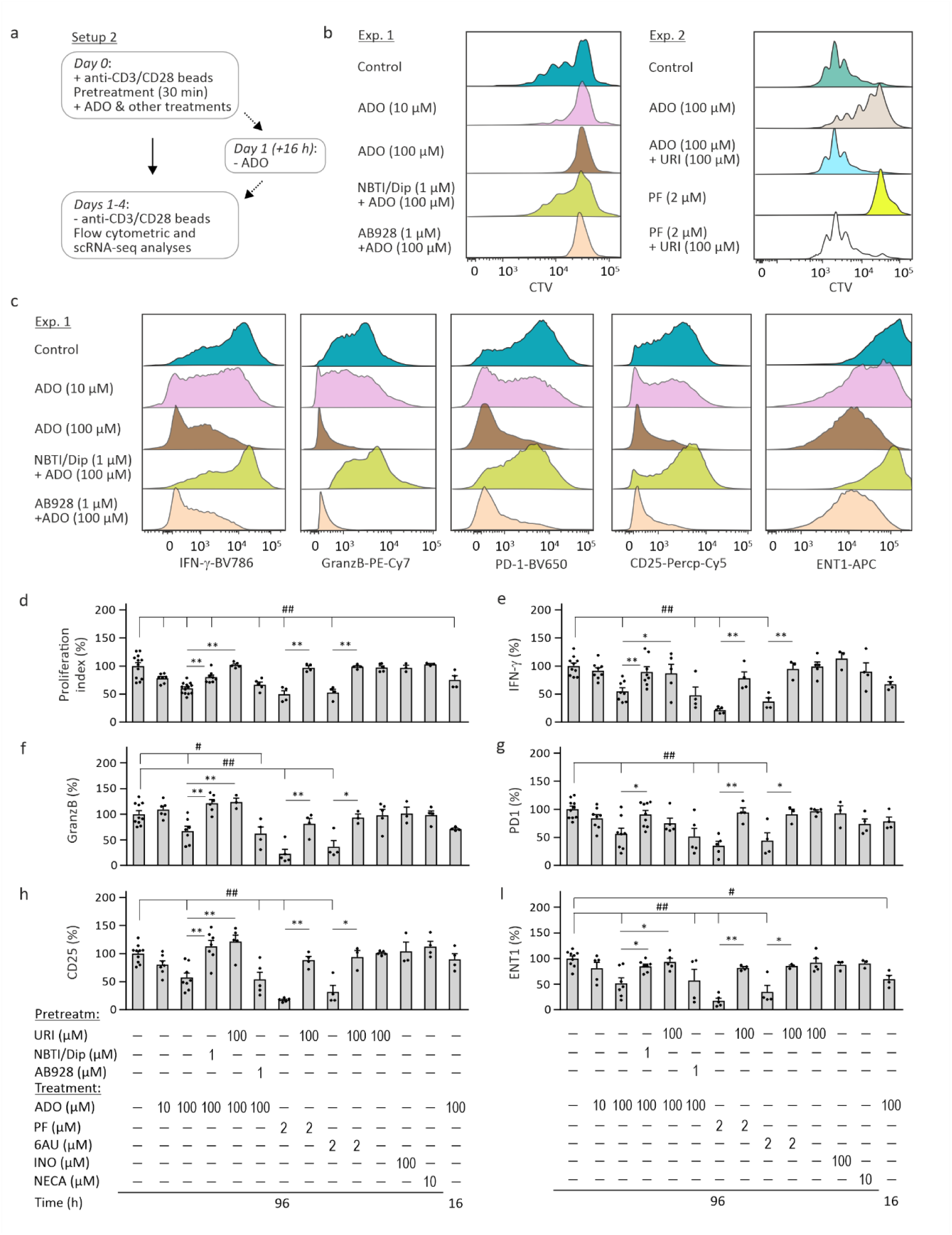
ADO suppresses transition of T cells from a quiescent to activated state via intracellular uptake through ENT1 and inhibition of pyrimidine metabolism. (a) Experimental setup to study the effect of ADO and other drugs on the proliferation and function of T cells during antigen stimulation. (b) T cells were labeled with a proliferation dye CTV, incubated for 96 hours with anti-CD3/CD28 beads and the indicated concentrations of ADO and other drugs, and analyzed by flow cytometry. Representative histograms of control and treated CTV-labeled T cells from two independent experiments are shown. (c) Flow cytometric analysis of the cytokine production and activation state of T cells. The graphs show the representative histograms for the production of IFN-γ and GranzB, and the expression of PD-1, CD25 and ENT1. Quantitative analysis of the proliferation index (d), synthesis of IFN-γ (e) and GranzB (f), and expression of PD-1 (g), CD25 (h), and ENT1 (i) in T cells activated with anti-CD3/CD28 beads and treated for 16-96 hours with ADO and other drugs. Mean fluorescence intensities were quantified using FlowJo and expressed as percentage of control (mean + SEM). Statistical significances were determined by one-way ANOVA with Dunnett’s multiple comparison test (^#^P<0.05 and ^##^P<0.01), or unpaired Student’s t-test (*P<0.05 and **P<0.01).

These findings corroborate the above data showing that ADO inhibits energy metabolism, triggers apoptosis and diminishes IFN-γ secretion by fully activated T cells, and extend these observations by demonstrating that ADO suppresses the transition of T cells from a quiescent to activated state. To determine whether the presence of ADO is required for sustained suppression of T cells, naïve T cells were incubated with anti-CD3/CD28 beads and 100 μM ADO for 16 hours, washed, and incubated with the beads for additional 80 hours (Fig. 5a). Even such short-term exposure to ADO partially affected the activation state of T cells, as evidenced by a decreased proliferation index (Fig. 5d) and low expression of ENT1 (Fig. 5i). However, the absence of significant changes in IFN-γ and GranzB production and CD25 and PD-1 expression suggests that ADO needs to be present in the assay medium throughout the entire period for efficient suppression of T cell activation (Fig. 5e-h).

### 6. scRNA-seq analysis reveals ADO-induced reprogramming in T cells during activation, characterized by impaired cell-cycle progression and type 1 interferon pathway activation

To investigate the mechanisms underlying the inhibitory effects of ADO during the late and early stages of T cell activation, we performed scRNA-seq analysis of T cells treated with anti-CD3/CD28 beads and ADO in two different ways. In the first setup, fully activated T cells were incubated for 24 hours without and with 100 μM ADO. Unbiased clustering identified distinct T cell clusters, with no significant changes in cluster composition observed between the control and ADO-treated cells (Supplementary Fig. 10). By contrast, when T cells were activated with anti-CD3/CD28 beads in the absence and presence of 100 μM ADO (setup 2), ADO markedly reshaped the T cell transcriptome, giving rise to distinct transcriptional clusters (Fig. 6). Changes in cluster composition were observed already at day 1: while control T cells were enriched in early activated *CD4^+^CD69*^+^ cells (cluster 1), *CD4^+^TAP1^+^* cells (cluster 3), and *CD4^+^TCF7*^+^ cells representing naïve CD4^+^ T cells (cluster 4), ADO-treated T cells preferentially accumulated in two CD4^+^ clusters characterized by the expression of *POLR2A* (clusters 7 and 8), while still retaining the naïve CD4^+^ T cells (cluster 4) (Fig. 6b, Supplementary Fig. 11). Following 3 days of activation, control T cells transitioned toward proliferative and activated states, including CD4^+^ clusters enriched for the cell-cycle-associated marker *CCNB1* (cluster 0), and genes encoding histones *H3C12, H4C4,* and *H3C8* (cluster 2), as well as CD8^+^ cluster enriched for vimentin, a marker associated with activation-related cytoskeletal remodeling (cluster 6) (Fig. 6a,b). However, ADO-treated cells retained a transcriptional profile resembling the early activation stage (clusters 1, 3, and 4). In addition, a distinct CD4^+^ cluster expressing a cell cycle regulator *EVI5* (cluster 5) emerged selectively in response to ADO treatment (Fig. 6a,b). Consistent with these findings, cell-cycle phase scoring based on canonical markers indicated that ADO treatment blocked the transition from G1 to S and G2/M phases at both day 1 and 3 (Supplementary Fig. 12).

**Fig. 6.**
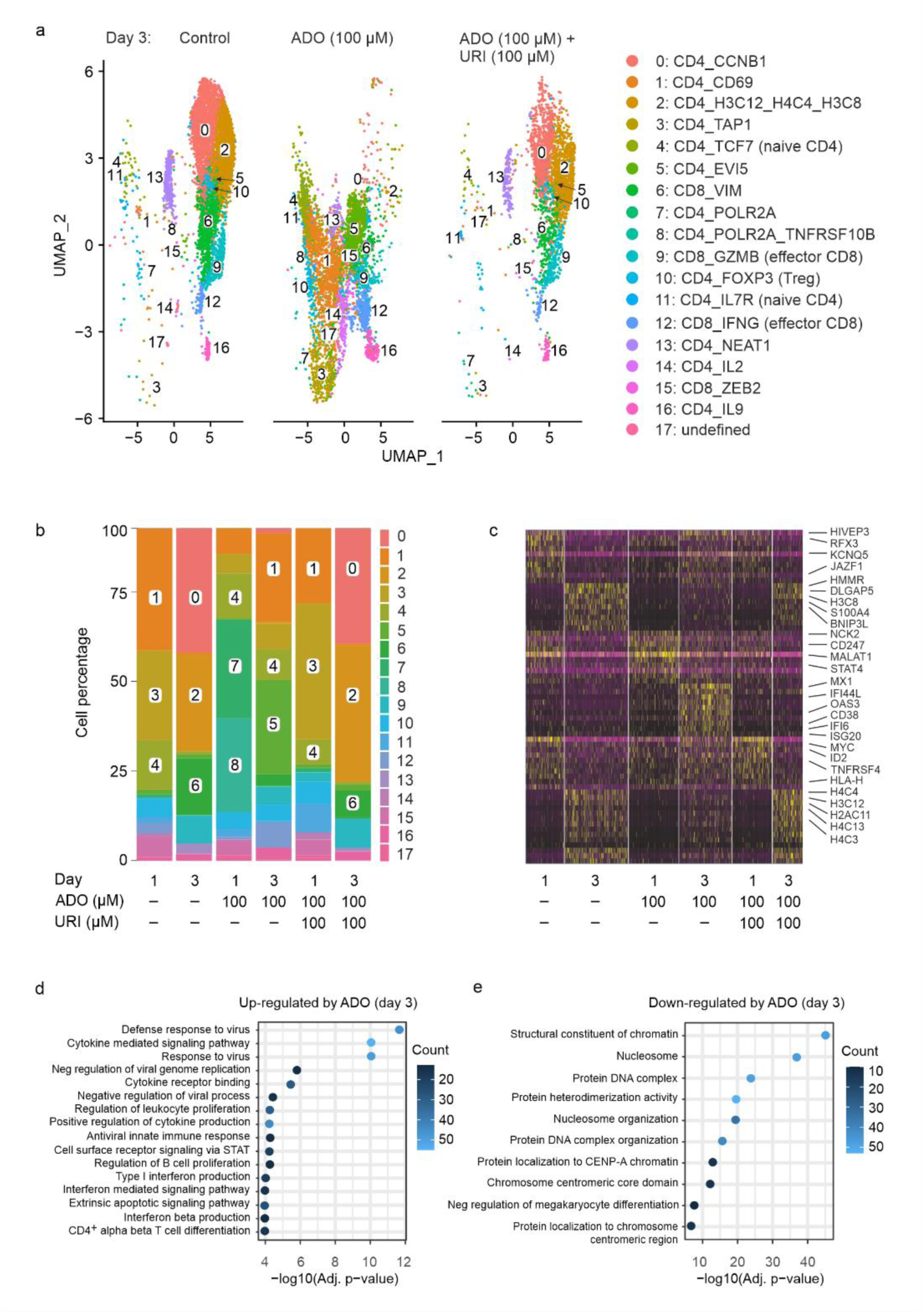
scRNA-seq analysis of human T cells activated with anti-CD3/CD28 beads in the presence of ADO and uridine. Peripheral blood T cells were isolated from two healthy adult volunteers and incubated for one or three days with anti-CD3/CD28 beads in the absence (control) and presence of 100 µM ADO and uridine (URI), as indicated. Transcriptome-based clustering of the single-cell expression profiles revealed 17 clusters, which were defined according to the expression of CD8, CD4 and other T cell markers (see Supplementary Fig. 11), as well as T cell cycle phase (S/G1/G2/M) (Supplementary Fig. 12). (a) UMAP plot of T cells activated for three days in the presence or absence of ADO and URI, color coded by cluster. (b) Stacked bar plot shows the percentage of cells assigned to each cluster across samples. (c) Heatmap illustrates the expression of the top differentially expressed genes between the control and treated T cells. The yellow color represents high expression, and purple represents low expression. Gene Ontology annotation enrichment analysis of the top genes up-regulated (d) and down-regulated (e) in ADO-treated T cells after three days of activation, determined using the EnrichR package. The color of the dots represents the number of enriched genes. FDR-corrected p values are shown.

Next, we performed a pseudo-bulk analysis of all T cells, irrespective of cluster identifications. This analysis revealed clear differences in transcriptional programs between control and ADO-treated T cells, which were already apparent at day 1 and became more pronounced following 3 days of activation (Fig. 6c). Control T cells displayed a strong proliferative and cell-cycle-associated transcriptional program, characterized by enrichment of mitotic regulators, such as *DLGAP5* and several histone genes, including *H3C8*, *H4C4*, and *H3C12*, consistent with active DNA replication and cell division. In contrast, ADO-treated cells were characterized by up-regulation of type 1 interferon-associated gene signatures, including antiviral effector genes *MX1*, *OAS3*, and *ISG20*, as well as interferon-responsive genes IFI44L and IFI6 (Fig. 6c). Additional Gene Ontology enrichment analysis of ADO-treated T cells revealed enrichment of pathways related to “Defense response to virus”, “Cytokine-mediated signaling pathway”, “Type I interferon response”, as well as other genes involved in cytokine signaling and antiviral immune responses, with these gene expression signatures emerging as early as day 1 (Supplementary Fig. 13a) and becoming increasingly robust by day 3 of treatment (Fig. 6d). At the same time, T cells activated in the presence of ADO for 1 (Supplementary Fig. 13b) and 3 (Fig. 6e) days exhibited significant down-regulation of genes associated with “DNA replication”, “Structural constituent of chromatin”, “Protein DNA complex”, “Nucleosome assembly”, and “Protein heterodimerization”. These findings demonstrate that ADO suppresses a transcriptional program already at early stages of T cell activation, which results in the impaired cell-cycle progression, reduced histone expression, and activation of type 1 interferon-responsive antiviral and stress-response pathways. Importantly, uridine supplementation restored the cluster composition of ADO-treated T cells to control levels, and reversed transcriptional changes induced by ADO (Fig. 6, Supplementary Figs. 11 and 12).

### 7. Molecular docking, molecular dynamics and free-energy analysis on human UMPS support direct binding of AMP to the OMPD active site

To understand the structural basis of inhibition of UMPS by ADO, we performed molecular dynamics (MD) simulations for binding of different substrates to OMPD, one of the two catalytic domains of this bifunctional enzyme responsible for catalytic conversion of orotate-derived OMP to UMP ^33, 34^ (Fig. 1a). All simulations used the human OMPD homodimer crystal structure (PDB: 2QCD, 2.03 Å resolution) as the reference ^34^. The OMP binding pose was obtained by superimposing PDB 2QCL (1.85 Å, D312N mutant) onto the wild-type frame (backbone RMSD = 0.26 Å). To assess whether other purines and pyrimidines can compete with OMP and UMP, candidate ligands were docked into the OMPD catalytic site using AutoDock Vina (Supplementary Table 1). Two structural features constrain ligand selection. First, the phosphate binding subsite formed by Lys281, Lys314 and backbone amides creates a positively charged cradle that accommodates one phosphate group. Therefore, polyphosphate tails (ATP, ADP, GTP), as well as ADO, which lacks this group entirely, cannot achieve strong binding. Second, the nucleobase pocket accommodates both pyrimidine (UMP, OMP) and purine (AMP) rings, but the hydrogen bonding pattern involving Asp312 and the backbone of Ser372 is optimized for the pyrimidine scaffold. Other nucleoside 5’-monophosphates GMP, CMP and IMP cannot be excluded by docking alone (Supplementary Table 1). Earlier competitive radiochemical enzymatic analyses of UMPS from mouse Ehrlich ascites cells ^35^ and human erythrocytes ^36^ revealed that UMP, AMP, CMP and IMP, but not GMP, inhibit OMPD activity of this bifunctional enzyme in the low micromolar range (10-100 µM). Consistent with these observations, recent isothermal titration calorimetry and high-resolution X-ray crystallographic studies of human UMPS demonstrated that CMP binds the OMPD active site with an affinity comparable to UMP, while GMP does not exhibit detectable binding ^33^. However, unlike the established link between ADO-driven AMP generation and T cell immunity, increasing intracellular CMP, IMP, or GMP levels through supplementation with the corresponding nucleosides, cytidine, inosine, and guanosine, had no impact on T cell function or metabolism in our study. Based on these considerations, subsequent detailed MD and free energy calculations were focused on UMP, OMP, and AMP.

Per-residue Molecular Mechanics Generalized Born Surface Area (MM-GBSA) decomposition (gmx_MMPBSA 1.6.4, igb = 5, mbondi3, last 50 ns, both sites combined, n = 6) identified Arg451, Lys314, Gln430, Lys281, Gly450, Tyr432, and Ser257 as the dominant active-site contributors and showed that the same residues anchor all three ligands. Arg451 forms the central salt bridge with the 5′-phosphate, Lys314 contacts the nucleobase, and Gln430/Lys281/Tyr432 line the pocket (Fig. 7a-d). Per-residue contributions are roughly twice as large for OMP at Arg451 as for AMP and UMP because OMP carries the additional C6-carboxylate charge (q = −3 versus −2 for AMP and UMP). Likewise, at Lys314 the OMP contribution exceeds that of AMP roughly two-fold, whereas the UMP contribution at this position is negligible (−0.2 ± 0.7 kcal/mol) (Fig. 7e). These are per-residue interaction contributions and do not by themselves rank the three ligands by total ΔG_bind, since the stronger OMP contacts are partly offset by larger desolvation penalty. Total MM-GBSA estimates overlap within their statistical uncertainty and are therefore not used to rank affinities (Fig. 7f). Across these trajectories UMP, OMP, and AMP all remained stably bound for the entire trajectory in every replicate. The nucleobase ring stayed close to its docked pose in all three ligands (heavy-atom RMSD 0.9-1.5 Å), whereas the 5′-phosphate sampled a larger conformational region (1.9-2.3 Å), consistent with the Arg451 salt bridge being the principal anchoring contact and the base providing the selectivity-determining contacts. Inter-site comparison gave a backbone RMSD of 1.4-1.6 Å, indicating the two binding sites of the homodimer support similar but not strictly identical conformations.

**Fig. 7.**
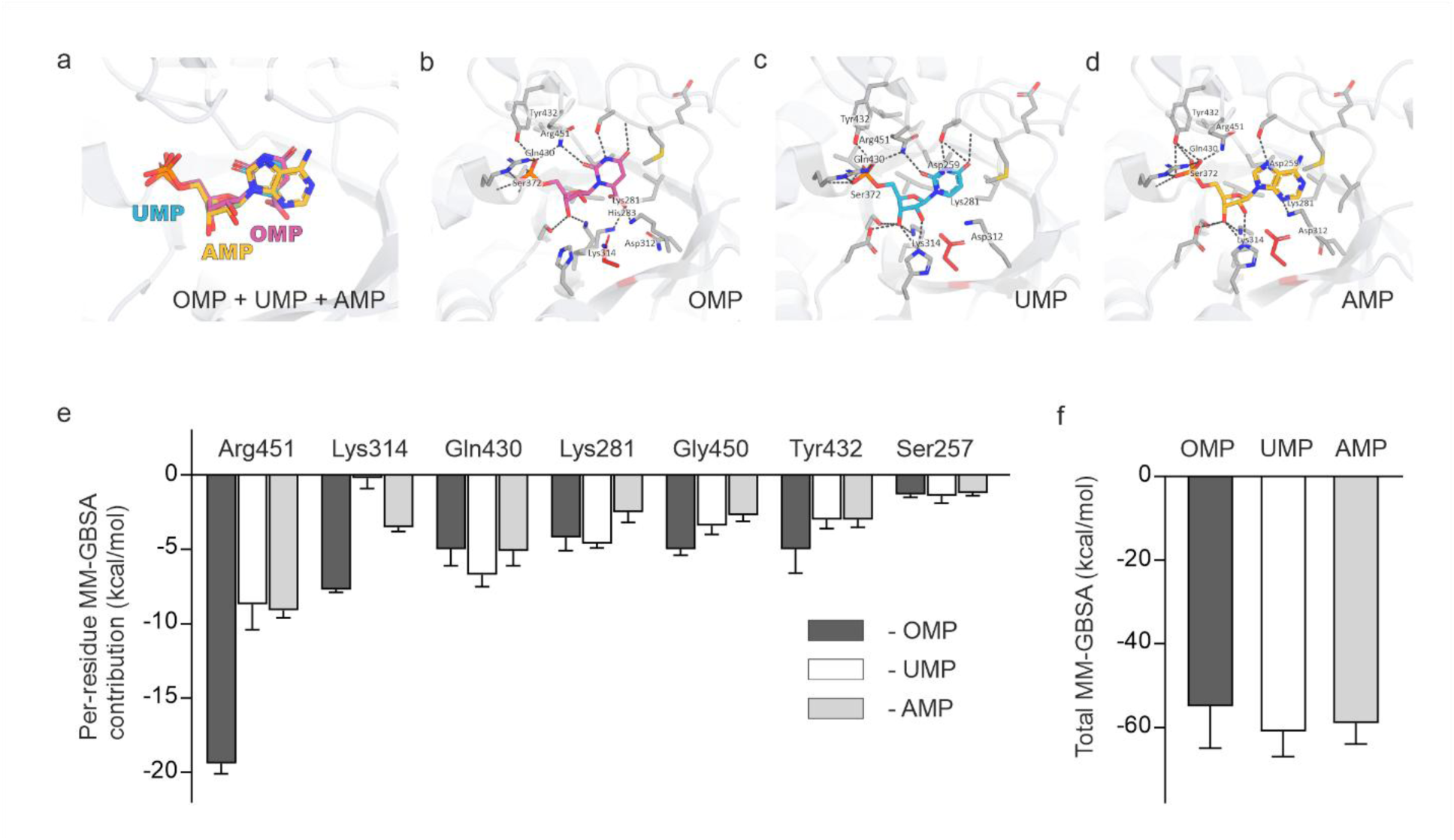
Binding modes and energetic analysis of UMP, OMP, and AMP at the human UMPS-OMPD active site. (a) Overlay of the three nucleotides in the OMPD pocket of chain A after superposition on 2QCD: UMP (cyan carbons, from 2QCD), OMP (pink carbons, from the 2QCL D312N mutant) and AMP (yellow carbons, modeled into 2QCD). All three ligands occupy a common volume, with their 5′-phosphate groups co-localized at the conserved anchor and their nucleobases diverging into the same aromatic-polar subsite. (b-d) Close-up views of OMP (b), UMP (c) and AMP (d) in the catalytic pocket. Pocket side chains within 4.5 Å of the ligand are drawn as gray sticks and labeled with full-length human UMPS numbering (UniProt P11172); the catalytic Asp312 is highlighted in red (this residue is Asn in the 2QCL mutant structure and is labeled Asp312 for cross-panel consistency); dashed lines indicate polar contacts within 3.5 Å. In all three complexes the 5′-phosphate is held by Arg451 and Lys314, while the base occupies the same subsite: the uracil O4/N3 face of UMP hydrogen-bonds to Gln430 and Tyr432, the C6-carboxylate of OMP engages Arg451 and Lys281, and the purine 6-amino group of AMP fills the position taken by the uracil C4 carbonyl. No co-crystal of AMP bound to human OMPD is available; the AMP pose shown is a model that was reproduced by independent AutoDock Vina redocking into 2QCD chain A. Net ligand charges at neutral pH: UMP −2, OMP −3 (C6-carboxylate), AMP −2. OMPD is catalytically active only as a dimer; one protomer is shown for clarity. The poses shown are the crystallographic and modeled reference structures, whereas the contact and energetic analyses reported in the text derive from the MD ensembles. (e) Per-residue MM-GBSA interaction contributions (kcal/mol, mean + SEM; n = 6, three MD replicates at each of the two protomer sites; final 50 ns of each trajectory) for the seven dominant active-site residues. OMP makes the strongest contacts, most markedly at Arg451 and Lys314, reflecting its additional C6-carboxylate charge. (f) Total MM-GBSA effective binding energies (mean + SEM). The three values overlap within their uncertainties, a known artifact of differencing large electrostatic and solvation terms for charged ligands and are therefore not used to rank affinities. The relative electrostatic binding component obtained from AMBER decharging thermodynamic integration, ΔΔG_dechg(AMP − UMP) = −1.68 ± 1.0 kcal/mol, indicates that AMP is as electrostatically complementary to the pocket as UMP.

The rigorous comparison currently available is an AMBER decharging thermodynamic-integration (TI) calculation of the electrostatic component of binding for AMP versus UMP, performed at the same active site with Boresch six-DOF restraints in the complex leg. The result, ΔΔG_dechg(AMP − UMP) = −1.68 ± 1.0 kcal/mol (electrostatic component only), indicates that the electrostatic complementarity of AMP to the OMPD active site is at least as favorable as that of UMP. Combined with stable pose retention over 100 ns of MD and the conserved Arg451 / Lys314 contact pattern, it indicates that AMP is sterically and electrostatically compatible with the catalytic pocket. Taken together, the MD, MM-GBSA contact pattern, and TI result are mutually consistent and provide structural and energetic support for the concept that ADO-derived AMP inhibits UMPS activity by direct binding at the OMPD active site.

## DISCUSSION

By investigating the combined features of cellular ADO metabolism and T cell immunity, we identified a link between these apparently interrelated processes. Major findings are summarized as follows: ADO triggers apoptosis, rewires energy metabolism, and impairs the effector functions of activated T cells, and also suppresses transition of T cells from a quiescent to an activated state. These effects of ADO can be phenocopied by inhibition of UMPS activity but reversed following pharmacological blockage of ENT1 or uridine supplementation. These data indicate that ADO inhibits T cell function through cellular uptake, sequential phosphorylation to ATP, and inhibition of pyrimidine nucleotide biosynthesis. This conclusion is reinforced by MD, MM-GBSA per-residue decomposition, and AMBER decharging TI on the human UMPS, suggesting that ADO metabolite AMP occupies the OMP/UMP catalytic pocket with the same Arg451 salt bridge to the 5′-phosphate and Lys314 contact to the nucleobase, exhibits complementarity to UMP, and is therefore sterically and electrostatically compatible with the OMPD active site.

This study identified purine metabolism in T cells as a dynamic network that undergoes reprogramming upon T cell activation via several interrelated mechanisms, including: (1) enhanced turnover of extracellular nucleotides through counteracting ATP-inactivating and ATP-regenerating pathways; (2) prevention of accumulation of extracellular ADO via down-regulation of CD73 and further elimination of residual ADO through high ecto-ADA activity and/or cellular uptake via ENT1; and (3) intracellular ADO metabolism through sequential phosphotransfer reactions. Additional scRNA-seq analysis of activated T cells revealed coordinated up-regulation of genes encoding key enzymes involved in the biosynthesis of purine and pyrimidine nucleotides, as well as ENT1. These findings support the current paradigm that activated T cells efficiently utilize extracellular ADO and other nucleosides, alongside sugars and amino acids, to meet their increased energetic and metabolic demands ^21, 23, 31^.

While the versatility of ADO as a potent signaling molecule is well recognized, the role of intracellular ADO metabolism in the control of T cell immunity is less characterized. Previous studies with *Ada^-/-^* mice ^37, 38^, *Adora2a^-/-^* mice with global or cre-mediated T cell deletion of A_2A_R ^39^, and human T cell lines and peripheral and tumor-infiltrating lymphocytes ^40, 41^ demonstrated the ability of ADO to suppress T cells via activation of A_2A_R and other ARs. In addition, the unique purinergic signature of resting T cells identified here, characterized by high expression of CD73 and A_2A_R together with low levels of ADA and ENT1, may create optimal conditions for maintaining T cell quiescence through enhanced production of extracellular ADO and continuous activation of A_2A_R. However, in the course of activation and metabolic rewiring, T cells become responsive to ADO, which influences their viability, bioenergetics and effector function mainly through intrinsic A_2A_R-independent mechanisms. In fact, pharmacological inhibition of ENT1 abolished all effects of ADO on T cells, whereas targeting the adenosinergic pathway using A_2A_R/A_2B_R agonists or antagonists had no detectable impact on any aspect of T cell function in our study. No changes in differentiation, viability and effector functions of T cells were detected after blocking ADA activity with EHNA or adding inosine instead of ADO. These findings contradict previous data showing that inosine supports T cell growth and function ^24^, promotes antitumor immunity via activation of Th1 cells ^42^, augments function and stemness of CAR-T cells ^43^, and activates energy expenditure in brown adipocytes ^44^, with most of these effects mediated by cellular inosine uptake via ENT1. This discrepancy can be resolved based on the following considerations. First, we used inosine at concentration of 100 µM, which is comparable with micromolar levels of inosine found in serum and interstitial milieu ^42, 45^. In contrast, previously reported effects of inosine on T cells were determined using supra-physiological 2 mM ^24, 42^ and up to 10 mM ^43^ concentrations of nucleoside. Second, contrary to temporary 2-3-fold ATP increase in ADO-treated activated T cells, inosine caused rather moderate ATP rise by ∼50%, which remained elevated for up to 24 hours, and not reversed by NBTI and dipyridamole (Fig. 3). These data suggest that ADO is primarily utilized through purine salvage pathway, whereas inosine is inactivated into hypoxanthine and ribose-1-phosphate followed by entry into the pentose phosphate pathway ^24^ (see Fig. 1a). Consistent with this conclusion are our radio-TLC data with [^3^H]ADO (Fig. 2d), and recent isotope-tracing mass spectrometry analysis of the biodistribution of [U-^15^N]ADO ^45^, showing that ADO is assimilated in different cells and tissues and directly phosphorylated via AMP to ATP without its prior metabolism to inosine. These data point to the different metabolic routes for ADO and inosine and their counteracting ability to suppress and promote T cell immunity, respectively.

ADO diverted ATP production in activated T cells from glycolysis and OxPhos toward energetically efficient purine salvage pathway (Fig. 3). At first sight, such metabolic rewiring is likely to be beneficial for T cells. In the nutrient-deprived TME, T cells face the challenge of competing with tumor and other adjacent cells for limited fuels, such as glucose, amino acids, and lipids ^21, 22, 46^. At the same time, cancer cells create an ADO-rich microenvironment through the enhanced release and metabolism of ATP ^8, 47, 48^, providing means for T cells to harness intratumoral ADO as an alternative energy source. However, concurrent inhibition of glycolysis may affect effector functions of T cells ^49, 50^, and reduce the amounts of vital precursor molecules that are essential for biosynthesis of nucleic acids, lipids, and amino acids ^23, 24^. Noteworthy, moderate and steady increases in intracellular ATP occurred in activated T cells treated with inosine or adenine, but they were not accompanied by shifts in T cell energetics and function. On the other hand, two UMPS inhibitors, PF and 6AU, phenocopied inhibitory effects of ADO without affecting total cellular ATP. These findings imply that other intermediate metabolite(s) of the ATP salvage chain mainly contribute to the suppression of T cell function, presumably by impeding de novo pyrimidine biosynthesis. The following evidence indicates that AMP is the most likely candidate mediating these inhibitory effects of ADO. The high-affinity and low-capacity enzyme ADK constitutes a major metabolic sink for intracellular ADO ^51^, and its expression is up-regulated upon T cell activation (Fig. 1f). TLC analysis confirmed that, following uptake by activated T cells, [^3^H]ADO is sequentially phosphorylated to [^3^H]ATP, with [^3^H]AMP accounting for ∼10% of the total radioactivity (Fig. 2d). Given that intracellular ATP in T cells is maintained in the millimolar range and further increases 2-3-fold upon ADO treatment (Fig. 3b), it can be assumed that AMP simultaneously reaches high micromolar levels - a concentration sufficient to inhibit UMPS activity ^35, 36^. Noteworthy, although direct interference of ADO with orotate production at the OPRT domain of UMPS has also been shown, this inhibition occurs only at millimolar concentrations of ADO ^35^. In line with this concept are our MD data suggesting that AMP likely occupies the OMPD catalytic pocket with the same anchoring contacts as natural substrate OMP and product UMP (Arg451 / 5′-phosphate salt bridge shared with OMP and UMP; Lys314 / base recognition shared with OMP) (Fig. 7). Collectively, the pose, contact pattern and electrostatic complementarity reported here establish that AMP can occupy the OMPD catalytic site of UMPS at the elevated intracellular concentrations reached in ADO-treated activated T cells, not that it is a tight-binding ligand of the isolated enzyme.

Our study provides new insights into the metabolic crosstalk between purine and pyrimidine pathways and their role in the fine-tuned regulation of T cell function. According to this scenario, ADO suppresses T cells via phosphorylation to ATP and inducing a concomitant imbalance in nucleotide and nucleoside levels. Although these transient shifts occur during the first hours of T cell exposure to ADO, previous studies have shown that T cells are particularly sensitive to perturbations in pyrimidine biosynthesis during a narrow time window corresponding to their first rounds of division ^52, 53^. Alterations in the AMP:ADP:ATP ratios may dictate the energy charge and effector functions of T cells in the blood and tissues ^31^, and modulate the activities of AMP-activated protein kinase (AMPK) ^14, 54^ and mechanistic target of rapamycin complex 1 (mTORC1) ^1, 55^ acting as important energy sensors and regulators of cellular metabolism. Besides its involvement in the salvage pathway, intracellular ADO may promote the accumulation of S-adenosylhomocysteine (SAH) through SAH hydrolase reaction, which in turn inhibits S-adenosylmethionine-dependent methyltransferases and reduces DNA and histone methylation ^51, 56^. Interestingly, depletion of cellular pyrimidines increases expression of antiviral genes through pathways involving Interferon Regulatory Factor 1 (IRF1), innate immune signaling and other stress responses ^57^. The latter observations fit well with our scRNA-seq data showing activation of interferon-responsive antiviral pathways in ADO-treated T cells, and concurrent down-regulation of genes involved in DNA packaging and nucleosome assembly, as well as histone genes controlling T cell proliferation.

Pyrimidine synthesis is an essential metabolic route in proliferating T cells, providing UTP and other nucleotides required for DNA replication, RNA synthesis, and cell differentiation ^1, 23, 53^. When de novo pathway is compromised, T cells adapt by salvaging uridine from the extracellular environment, enabling the maintenance of clonal expansion, viability, and metabolic activity ^52, 53, 58^. Indeed, disruption of de novo pyrimidine synthesis through genome-editing approaches or enzyme inhibitors prevented T cell differentiation ^51, 52^, triggered T cell death ^58^, impaired T cell bioenergetics, viability and cytotoxicity (present study), and induced auxotrophy in human cell lines and pluripotent stem cells ^58^, with all these adverse effects being rescued by uridine supplementation. These findings, together with evidence that uridine counteracts the suppressive and cytotoxic effects of ADO on T cells by bypassing AMP-mediated inhibition of UMPS ^8^ (present study), may pave the way for new combination therapy aimed at restoring impaired T cell functions. Uridine is the most abundant circulating nucleoside, with plasma concentrations in humans ranging from 3 to 8 μM ^59^. A recent randomized pilot study in patients with asymptomatic bacteriuria showed that adjuvant oral supplementation with uridine and pyruvate protects the proliferative capacity of effector T lymphocytes from mitotoxic antibiotics ^60^. The FDA-approved uridine prodrug uridine triacetate (Vistogard®), currently used to treat severe 5-fluorouracil- and capecitabine-induced toxicity, exhibits better tolerability than uridine and markedly elevates uridine concentrations in human serum ^61^. Therefore, novel therapeutic strategies may include uridine-based therapeutic interventions, in combination with ENT1 blockers, and small-molecule inhibitors or antibodies directed against CD39 and CD73, with the desired outcome to reduce generation of ADO in the TME and limit its intracellular supply and adverse effects on pyrimidine metabolism. To achieve maximal inhibition of ADO transport, we treated T cells with NBTI acting as a potent inhibitor of human ENT1 ^29, 62^, in combination with a pleiotropic drug dipyridamole mediating diverse effects through the inhibition of cellular ADO uptake via ENT1 and/or phosphodiesterase activity ^13, 30^. Interestingly, while the later compound is widely used as a platelet aggregation inhibitor, it has the potential to be repurposed as an anti-tumor drug against lymphoid neoplasms and other malignancies ^63^. These findings provide rationale for revisiting the current view of biological actions of other FDA-approved drugs inhibiting ENT1, such as P2Y_12_ receptor antagonist ticagrelor acting as antithrombotic drug ^13^, anti-cancer drugs clofarabine and gemcitabine, anti-viral drugs abacavir, fialuridine, and remdesivir ^62^, and the analgesic and vasodilatory drug dilazep ^12^.

In conclusion, our study provides insights into mechanisms underlying the inhibitory effects of ADO on T cell immunity. Mechanistically, ADO mediates its effects through cellular uptake via ENT1, ADK-mediated phosphorylation to AMP, and further to ADP and ATP, and AMP-induced inhibition of the OMPD catalytic site of UMPS. Elucidating the precise mechanisms by which uridine alleviates the immunosuppressive effects of ADO and exploring the therapeutic potential of purine and pyrimidine metabolic pathways in cancer immunotherapy will be important areas of future research.

## MATERIALS AND METHODS

### Reagents and antibodies

Detailed information on primary antibodies used in this study is provided in Supplementary Table 2. Secondary Alexa Fluor® 488-, 546-, 633- and 750-conjugated goat anti-mouse, anti-rabbit and anti-chicken immunoglobulins, Alexa Fluor® 546-conjugated Phalloidin, ProLong® Gold Antifade reagent with 4,6-diamidino-2-phenylindole (DAPI), and GIBCO OpTmizer™ CTS™ T Cell Expansion Serum Free Medium (SFM) were from ThermoFisher Life Technologies. Recombinant human interleukin-2 “Improved Sequence” (IL-2 IS) and interleukin-7 (IL-7) were from Miltenyi Biotec. Cy3™-conjugated donkey anti-guinea pig IgG was from Jackson Immuno Research Laboratories (West Grove, PA). AB928 (also known as etrumadenant) was from Cayman Chemical (Michigan, USA). [2,8-^3^H]ADP, [2,8-^3^H]AMP and [2,8-^3^H]ADO were from American Radiolabeled Chemicals Inc. (Campro Scientific, The Netherlands). Benzylacyclouridine (BAU) and Compound 20874830-2 (C2087) were from MedChemExpress®. Dipyridamole, *S*-(4-nitrobenzyl)-6-thioinosine (NBTI, also known as NBMPR), 5’-(N-ethylcarboxamido)adenosine (NECA), P^1^,P^5^-di(adenosine-5’)-pentaphosphate (Ap_5_A), erythro-9-(2-hydroxy-3-nonyl)adenine (EHNA), 2-Imino-1-imidazolidinineacetic acid (cyclocreatine, cCr), 4-hydroxy-5-β-D-ribofuranosyl-1*H*-pyrazole-3-carboxamide (pyrazofurin, PF), 6-azauridine (6AU), ADO, other nucleosides, nucleotides and adenine, and other chemicals were from Sigma-Aldrich.

### T cell isolation, stimulation and treatment

Healthy volunteers between the ages of 24 and 62 participated in the study. The procedure of blood collection and T cell isolation was reviewed by the local Ethics Committee of the University of Turku and approved by the Provincial State Office of Western Finland with the license ETMK 43/1801/2015. All participants were required to sign a written consent form. T cells were isolated from peripheral blood using the EasySep™ T cell Isolation Kit (STEMCELL Technologies), according to the manufacturer’s instructions. In some experiments, T cells were additionally separated into CD4^+^ helper T cells and CD8^+^ effector T cells by immunomagnetic selection using the EasyStep™ Human Naïve CD8^+^ T Cell Isolation Kit II (Stem Cell). T cells were resuspended in complete T cell medium (comprising: OpTmizer™ CTS™ T Cell Expansion SFM, 2 mM L-glutamine, 100 U/ml penicillin, and 100 µg/ml streptomycin), maintained at 37°C in a humidified incubator with 21% O_2_ and 5% CO_2_, and subsequently activated and subjected to various treatments in the following ways: In the first approach, T cells were divided into two groups. One part was activated by incubating the cells for four days in complete T cell medium containing 30 U/ml IL-2 IS and human T-Activator CD3/CD28 Dynabeads™ (25 µl per 10^6^ cells, ThermoFisher). Thereafter, the beads were removed and activated T cells resuspended in T cell medium containing IL-2 IS. Another part of the cells was maintained in a resting state by culturing in T cell medium containing IL-7 (10 ng/ml) instead of IL-2 IS. Activated and resting T cells were incubated for up to 24 hours with ADO (0-100 µM) and other drugs, as outlined in Figure 3a. In the second experimental setup, ADO and other treatments were added to T cells within the first two hours of activation with anti-CD3/CD28 Dynabeads™, and then either washed out after 16 hours or kept in the assay medium for the entire activation period (Figure 5a).

### TLC analysis of extracellular and intracellular purine metabolism

Human T cells were added to 96-well flat bottom clear plates at a density of 120000 cells per well. Ectoenzymatic activities were determined by incubating T cells with ^3^H-labeled and unlabeled substrates in a final volume of 100 μl T cell medium supplemented with 5 mM β-glycerophosphate in the following ways: (i) for NTPDase1/CD39 assay, the cells were incubated for 60 min with 50 μM [^3^H]ADP in the presence of 100 μM Ap_5_A (for blocking [^3^H]ADP transphosphorylation through reverse AK reaction); (ii) ecto-5’-nucleotidase/CD73 was assayed by incubating T cells for 60 min with 50 μM [^3^H]AMP; (iii) AK and NDPK activities were measured by incubating T cells for 45 min with 500 μM γ-phosphate-donating ATP and 200 μM [^3^H]AMP or 200 µM [^3^H]ADP as respective enzyme substrates; (iv) ADA was assayed by incubating the cells for 60 min with 20 μM [^3^H]ADO. To measure cellular ADO uptake and its intracellular metabolism, T cells were incubated for 60 min with 10 μM [^3^H]ADO in a final volume of 200 μl T cell medium, washed with PBS to remove non-bound radioactivity, and lysed in the appropriate cell lysis buffer (Perkin Elmer). In some competitive assays, T cells were pretreated with 10 µM EHNA or a mixture of NBTI and dipyridamole (final concentration 1 μM for each) for 30 min prior to addition of [^3^H]ADO. Aliquots of the mixture were applied onto Alugram® SIL G/UV_254_ TLC sheets (Machery-Nagel, Duren, Germany), separated by TLC, and then either quantified for the amounts of ^3^H-labeled substrates and their metabolites or exposed to Kodak BioMax MS film (Carestream, Rochester, NY) for autoradiographic imaging, as described elsewhere ^15, 28^.

### Flow cytometry

T cells were activated and treated as described above, washed with PBS containing 2% fetal calf serum (FCS) (staining buffer), and subsequently incubated with Human TruStain FcXTM (BioLegend) to block non-specific Fc-receptor binding. Next, cells were stained with primary fluorochrome-conjugated antibodies (2-5 µg/ml diluted in staining buffer). For intracellular staining, T cells were fixed and permeabilized using the Cytofix/Cytoperm™ Fixation/Permeabilization Solution Kit (BD Biosciences), following incubation with the respective antibodies (2-5 µg/ml diluted in permeabilization buffer). For cytokine staining, T cells were pre-treated with Brefeldin A and Cell Activation Cocktail (BioLegend) for 4 hours prior to FACS staining. T cell proliferation was evaluated by labeling freshly isolated T cells with CellTrace™ Violet (CTV, ThermoFisher Scientific), according to the manufacturer’s instructions. Staining of apoptotic and dead cells was performed using the APC Annexin V Apoptosis Detection Kit with 7AAD (BioLegend). Flow cytometry data were acquired using an LSR Fortessa flow cytometer (BD Biosciences) and analyzed using FlowJo-v10.10.0 software (FlowJo LLC).

### Immunofluorescence staining

T cells were added onto poly-D-lysine-coated 13-mm coverslips in 24-well plates (∼3 × 10^5^ cells/well) and incubated for 2 hours in T cell medium. The wells were washed with PBS to remove non-adherent cells. The adherent T cells were fixed for 5 min with PBS containing 4% paraformaldehyde and additionally incubated for 3 min with 0.2% Triton X-100. The fixed and permeabilized T cells were processed for staining of membrane-bound nucleotidases, ENT1, and intracellular filamentous and cytoskeletal structures, as described earlier ^15^. The coverslips were mounted with ProLong® medium with DAPI. The images were examined using a 3i spinning disk confocal microscope CSU-W1 with Plan-Apochromat 100×/1.4 objective (Carl Zeiss) and Slidebook 6.0 software (Intelligent Imaging Innovations, Inc.). Z-stacks of the resting and activated T cells were captured at identical exposure times for each antibody and further acquired in parallel by using Imaris 8.4 (Bitplane) and Adobe Photoshop CS6.

### Measurement of intracellular ATP

Resting and activated T cells were added to a 96-well clear U-shaped plate at a density of 200000 and 100000 cells per well, respectively. The cells were pre-treated for 30 minutes at 37°C in a final volume of 200 μl complete T cell medium containing ENT1 inhibitors NBTI and dipyridamole (1 µM each), ADA inhibitor EHNA (10 µM), the inhibitors of UMP synthase PF (2 μM) and 6AU (10 μM), the inhibitor of creatine kinase cCr (200 μM), or vehicle containing 0.01% DMSO (control). Different concentrations of ADO, other nucleosides and adenine were added to the cells, and incubated for different time intervals. At the end of treatment, T cells were washed two times with 300 µl PBS, lysed using the appropriate cell lysis buffer, and assayed for intracellular ATP by using luciferin-luciferase kit ATPLite (Revvity Health Sciences Inc., ThermoFisher). Protein concentration in T cell lysates was determined spectrophotometrically using the BCA Protein Assay Kit (Pierce, Rockford, IL). All measurements were performed with Tecan Infinite M200 microplate reader (Salzburg, Austria).

### T cell metabolic assay

T cell metabolism was evaluated using the Agilent Seahorse XF Cell Mito Stress Kit, according to the manufacturer’s guidelines. Briefly, one day prior to the experiment, the XFe96/XF Pro Sensor Cartridge was hydrated at 37°C in a non-CO_2_ incubator. The Seahorse XF PDL cell culture plate was coated with 50 µl Poly-L-lysine solution (Sigma-Aldrich) for 5 minutes. The plate was then washed twice with H_2_O, allowed to air-dry, and kept at 37°C in a non-CO_2_ incubator. On the day of the experiment, T cells, treated as specified above, were harvested, washed in XF RPMI Medium supplemented with 10 mM glucose, 1 mM pyruvate, and 2 mM glutamine (Agilent), and resuspended in the same buffer at a concentration of 2×10^6^ cells per ml. Next, 50 µl of T cell suspension was added to the pre-coated XF cell culture microplate, and metabolic profiling was conducted using the Seahorse XFe96 Analyzer. The results were analyzed using Agilent’s web-based software Seahorse Analytics.

### scRNA-seq analysis of DC-primed T cells

DCs were isolated from non-tumor draining lymph nodes obtained from two breast cancer patients undergoing mastectomy and axillary lymph node evacuation at the Turku University Hospital, as previously described ^64^. The collection was done under the license ETMK: 132/2016. All patients gave written informed consent prior to sample collection, and the study was conducted in accordance with the Declaration of Helsinki. Obtained single-cell suspensions were incubated with TruStain FcX, and stained with AF488 anti-CD20, AF488 anti-CD3, BV421 anti-HLA-DR, PE anti-CD141, PerCP-Cy5.5 anti-CD11C, PE-Cy7 anti-CD1C. CD3^-^CD20^-^HLA-DR^+^CD11C^+^CD141^-^CD1C^+^ DCs were sorted using SH800S sorter (Sony) into RPMI-1640 medium containing 10% FCS. 5000 DCs were co-cultured with 50000 allogenic naïve T cells from the venous blood of two healthy donors, isolated as described above. Six days later, DCs and T cells were collected, counted, and processed for scRNA-seq, following 10x Genomics guidelines. scRNA-seq libraries were prepared according to the manufacturer’s instructions using the Chromium Single Cell 3’ Library and Gel Bead Kit v3 (10x Genomics) along with the Chromium Single Cell A Chip Kit. Sequencing was performed using the Illumina NovaSeq platform. Each sample was sequenced to achieve an average depth of 50000 reads per cell. Postprocessing, including demultiplexing, read alignment, and quality control was performed at the Finnish Functional Genomics Centre of the University of Turku using the 10x Genomics Cell Ranger package (ver. 5.0.1).

### scRNA-seq analysis of ADO-treated T cells

T cells were isolated from the blood of healthy donors and treated in two different ways. In the first setup, T cells from two donors were stimulated for four days with anti-CD3/CD28 Dynabeads™ and additionally incubated for 24 hours without and with 100 μM ADO. In another experimental setup, freshly isolated T cells were activated with anti-CD3/CD28 beads in the absence and presence of 100 μM ADO and/or uridine. Cells were then processed using Evercode^TM^ fixation v2 kit and Evercode^TM^ WT v2 kits from Parse Biosciences, according to the manufacturer’s instructions. The eight resulting sub-libraries were pooled and sequenced on an Illumina Novaseq sequencer (S2 flowcell). For data processing, the Parse Bioscience processing pipeline (v1.1.1) was used with default settings to align sequencing reads to GRCh38 human genome and to demultiplex samples. Each of the eight sub-libraries was first processed individually using the command split-pipe --mode all, and the output of the eight sub-libraries was combined using the command split-pipe --mode combine.

### scRNA-seq data processing and clustering

Downstream processing was performed using the Seurat R package v5. For quality control, we removed genes which were expressed in less than 5-10 cells. We also filtered out the cells with less than 200 -500 genes and cells with greater than 7500-15000 genes and 15-25% mitochondrial genes. Data were normalized by the function of NormalizeData, variable genes across the single cells were detected by FindVariableFeatures function, and the data were scaled by ScaleData function. For principal component analysis (PCA), either reciprocal principal component analysis (RPCA) or conventional PCA was used. RPCA was used to remove technical batch effects between patient samples in the dataset of DC-primed T cells. Then, a graph-based clustering approach was used by running FindNeighbors and FindClusters functions. The clustering was visualized with Uniform Manifold Approximation and Projection (UMAP). Differentially expressed genes between the clusters and treatments were calculated with the functions FindMarkers or FindAllMarkers. Pathway enrichment analysis of DEGs was performed using Enricher. For pseudobulk DEG analysis, clusters were separated into non-treated or treated T cells and DEGs between these groups were analyzed by using FindAllMarkers. Cell-cycle scoring was performed using CellCycleScoring in Seurat.

### Data availability

scRNA-seq data have been deposited in the GEO database under accession number GEO: GSExxxx.

### scRNA-seq analysis of publicly available T cell dataset

We utilized publicly available scRNAseq data of human T cells ^26^. In this dataset, T cells were isolated from the blood of two healthy donors and cultured for 16 h in the media alone or in the presence of anti-CD3/anti-CD28 antibodies. The data have been deposited at the National Center for Biotechnology Information (NCBI) Gene Expression Omnibus (GEO) database with accession number GSE126030. The data were analyzed using Seurat v5. Low-quality cells were filtered out as described above. Dataset from two donors were integrated using RPCA and analyzed using a graph-based clustering approach.

### Molecular docking, molecular dynamics and free-energy calculations

The human UMPS OMPD domain was taken from the homodimeric structure PDB 2QCD (2.03 Å)^34^, and the reference OMP pose from the D312N mutant 2QCL (1.85 Å) superimposed on it (backbone RMSD 0.26 Å). The compounds listed in Supplementary Table 1 were docked into the catalytic site with AutoDock Vina 1.2.0 ^65^ (exhaustiveness 32, 25 Å box centered on the UMP 5′-phosphate subsite) and ranked by predicted affinity; AMP, UMP and OMP were retained for simulation. Each retained ligand was placed in both active sites of the pH 7-protonated 2QCD homodimer, parameterized with AMBER ff14SB (protein) ^66^ and GAFF ^67^ with AM1-BCC ^68^ charges (ligand; net charge −2 for AMP and UMP, −3 for OMP), solvated in a cubic TIP3P box (1.0 nm padding, 0.15 M NaCl, ∼99,000 atoms), energy-minimized, equilibrated under position restraints (NVT 100 ps, NPT 2 × 100 ps) and simulated unrestrained for 100 ns in GROMACS 2025.4 ^69^ at 298.15 K and 1 bar (V-rescale and C-rescale coupling, PME, 1.0 nm cut-offs, LINCS constraints, hydrogen mass repartitioning, 4 fs time step). Three replicates per ligand gave n = 6 trajectories (three replicates × two protomer sites). From the final 50 ns of each trajectory, effective binding energies and per-residue decomposition (idecomp = 2, 6 Å shell) were obtained by single-trajectory MM-GBSA (gmx_MMPBSA 1.6.4; igb = 5, mbondi3 radii, 0.15 M salt) ^70^, and ligand RMSD and per-residue contact frequencies with MDAnalysis after a Kabsch fit on the 19 pocket residues. Values are reported as mean ± SEM over the six trajectories, which share one homodimer and are therefore not strictly independent, so the standard errors are a lower bound. The electrostatic component of AMP versus UMP binding was compared by AMBER decharging thermodynamic integration (dual-topology soft core, 12 λ windows of 2 ns, Boresch six-degree-of-freedom restraints in the complex leg, trapezoidal integration of the ensemble-averaged ∂U/∂λ. Residues are numbered as in full-length human UMPS (UniProt P11172).

### Statistical analysis

Each experiment was conducted using at least two technical replicates for all treatments. Unless otherwise stated, results are expressed as scatter plots with bars (mean + SEM), with each symbol displaying independent experiment performed with T cells isolated from one donor. Comparisons between two groups were performed using Student’s t test. The statistical significance levels were denoted as *P<0.05 and **P<0.01. Multiple groups were compared using one-way analysis of variance (ANOVA) with Dunnett’s multiple comparison test. ^#^P<0.05 and ^##^P<0.01 were considered statistically significant. Statistical tests corresponding to each experiment are indicated in the figure legends. All results were analyzed with GraphPad Prism 7.5 software (GraphPad, San Diego, CA).

## Acknowledgments

We thank Dr. Jouko Sandholm at the Cell Imaging and Cytometry Unit of Turku Bioscience Centre for microscopy support and Biocenter Finland for microscopy and flow cytometry equipment. We are also grateful to Prof. Jean Sevigny (Université Laval, QC, Canada) for providing the antibodies against human CD73 and CD39, and Dr. Pia Boström and Prof. Ilkka Koskivuo (Turku University Hospital, Finland) for providing human lymph node samples for scRNA-seq analysis. We acknowledge support from the Research Council of Finland, the Finnish Cancer Foundation, Sigrid Juselius Foundation, and Jane and Aatos Erkko Foundation. This article is based upon work from PRESTO COST Action CA21130, supported by COST (European Cooperation in Science and Technology). Financial support of UJVD, JSB, NK, and CW by the Erasmus+ EU Traineeship Program is greatly acknowledged.

## Author contributions

Conceptualization and supervision: AT, GGY; Methodology and investigation: KL, UJVD, NK, SS, JSB, CW, AT, GGY; Visualization: KL, GGY; Molecular docking: AZ, NJT, OG; Funding acquisition: SJ, AT; Manuscript writing, review, and editing: KL, SJ, AT, SEM, GGY.

## References

1. Ali ES, Ben-Sahra I. Regulation of nucleotide metabolism in cancers and immune disorders. Trends in Cell Biology, (2023).

2. Mullen NJ, Singh PK. Nucleotide metabolism: a pan-cancer metabolic dependency. Nat Rev Cancer 23, 275–294 (2023).

3. Ralevic V, Burnstock G. Receptors for purines and pyrimidines. Pharmacol Rev 50, 413–492 (1998).

4. Vijayan D, Young A, Teng MWL, Smyth MJ. Targeting immunosuppressive adenosine in cancer. Nat Rev Cancer 17, 709–724 (2017).

5. IJzerman AP, Jacobson KA, Müller CE, Cronstein BN, Cunha RA. International Union of Basic and Clinical Pharmacology. CXII: Adenosine Receptors: A Further Update. Pharmacological Reviews 74, 340–372 (2022).

6. Allard B, Allard D, Buisseret L, Stagg J. The adenosine pathway in immuno-oncology. Nat Rev Clin Oncol 17, 611–629 (2020).

7. Yegutkin GG, Boison D. ATP and Adenosine Metabolism in Cancer: Exploitation for Therapeutic Gain. Pharmacological Reviews 74, 797–822 (2022).

8. Sanders TJ, et al. Inhibition of ENT1 relieves intracellular adenosine-mediated T cell suppression in cancer. Nat Immunol 26, 854–865 (2025).

9. Lu Q, Newton J, Hsiao V, Shamirian P, Blackburn MR, Pedroza M. Sustained adenosine exposure causes lung endothelial barrier dysfunction via nucleoside transporter-mediated signaling. Am J Respir Cell Mol Biol 47, 604–613 (2012).

10. Xu Y, et al. Regulation of endothelial intracellular adenosine via adenosine kinase epigenetically modulates vascular inflammation. Nat Commun 8, 943 (2017).

11. Mikdar M, Serra M, Azouzi S. Adenosine signaling in promoting the balance between erythropoiesis and myelopoiesis. Curr Opin Hematol 32, 199–205 (2025).

12. Wright NJ, et al. Design of an equilibrative nucleoside transporter subtype 1 inhibitor for pain relief. Nat Commun 15, 10738 (2024).

13. Cattaneo M, Schulz R, Nylander S. Adenosine-mediated effects of ticagrelor: evidence and potential clinical relevance. Journal of American College of Cardiology 63, 2503–2509 (2014).

14. Virtanen SS, et al. Adenosine Inhibits Tumor Cell Invasion via Receptor-Independent Mechanisms. Mol Cancer Res 12, 1863–1874 (2014).

15. Losenkova K, et al. Compartmentalization of adenosine metabolism in cancer cells and its modulation during acute hypoxia. Journal of Cell Science 133, jcs241463 (2020).

16. Ichikawa A, et al. Effect of adenosine and adenosine 5’-monophosphate on cell division of cultured mastocytoma P-815 cells. J Pharmacobiodyn 3, 123–135 (1980).

17. Festag J, et al. Preventing ATP Degradation by ASO-Mediated Knockdown of CD39 and CD73 Results in A2aR-Independent Rescue of T Cell Proliferation. Mol Ther Nucleic Acids 21, 656–669 (2020).

18. Marius W, et al. Expression of CD39 is associated with T cell exhaustion in ovarian cancer and its blockade reverts T cell dysfunction. Oncoimmunology 13, 2346359 (2024).

19. Bergero G, et al. Purinergic signaling modulates CD4+ T cells with cytotoxic potential during Trypanosoma cruzi infection. J Clin Invest 135, (2025).

20. Allard D, et al. Adenosine Uptake through the Nucleoside Transporter ENT1 Suppresses Antitumor Immunity and T-cell Pyrimidine Synthesis. Cancer Res 85, 692–703 (2025).

21. Chapman NM, Boothby MR, Chi H. Metabolic coordination of T cell quiescence and activation. Nat Rev Immunol 20, 55–70 (2020).

22. Reina-Campos M, Scharping NE, Goldrath AW. CD8(+) T cell metabolism in infection and cancer. Nat Rev Immunol 21, 718–738 (2021).

23. Zhu J, Thompson CB. Metabolic regulation of cell growth and proliferation. Nat Rev Mol Cell Biol 20, 436–450 (2019).

24. Wang T, et al. Inosine is an alternative carbon source for CD8(+)-T-cell function under glucose restriction. Nat Metab 2, 635–647 (2020).

25. Skinner OS, et al. Salvage of ribose from uridine or RNA supports glycolysis in nutrient-limited conditions. Nat Metab 5, 765–776 (2023).

26. Szabo PA, et al. Single-cell transcriptomics of human T cells reveals tissue and activation signatures in health and disease. Nat Commun 10, 4706 (2019).

27. Schneider E, et al. CD73-mediated adenosine production by CD8 T cell-derived extracellular vesicles constitutes an intrinsic mechanism of immune suppression. Nat Commun 12, 5911 (2021).

28. Yegutkin GG. Enzymes involved in metabolism of extracellular nucleotides and nucleosides: Functional implications and measurement of activities. Crit Rev Biochem Mol Biol 49, 473–497 (2014).

29. Boswell-Casteel RC, Hays FA. Equilibrative nucleoside transporters-A review. Nucleosides Nucleotides Nucleic Acids 36, 7–30 (2017).

30. Balakumar P, et al. Classical and pleiotropic actions of dipyridamole: Not enough light to illuminate the dark tunnel? Pharmacol Res 87, 144–150 (2014).

31. Chikina AS, Corre B, Lemaitre F, Bousso P. Spatiotemporal regulation of energetic charge dictates T cell function. Nat Commun 17, 770 (2026).

32. Di Biase S, et al. Creatine uptake regulates CD8 T cell antitumor immunity. J Exp Med 216, 2869–2882 (2019).

33. Kirck LL, Santagostino E, Brandhoff L, Simeth NA, Tittmann K. Expected and Unexpected “Guests” at the Active Site of Human Orotidine 5’-Monophosphate Decarboxylase. Biochemistry 64, 4542–4554 (2025).

34. Wittmann JG, Heinrich D, Gasow K, Frey A, Diederichsen U, Rudolph MG. Structures of the human orotidine-5’-monophosphate decarboxylase support a covalent mechanism and provide a framework for drug design. Structure 16, 82–92 (2008).

35. Traut TW, Jones ME. Inhibitors of orotate phosphoribosyl-transferase and orotidine-5’-phosphate decarboxylase from mouse Ehrlich ascites cells: a procedure for analyzing the inhibition of a multi-enzyme complex. Biochem Pharmacol 26, 2291–2296 (1977).

36. Tax WJ, Veerkamp JH. Inhibition of orotate phosphoribosyltransferase and orotidine-5’-phosphate decarboxylase of human erythrocytes by purine and pyrimidine nucleotides. Biochem Pharmacol 28, 829–831 (1979).

37. Apasov S, Chen JF, Smith P, Sitkovsky M. A(2A) receptor dependent and A(2A) receptor independent effects of extracellular adenosine on murine thymocytes in conditions of adenosine deaminase deficiency. Blood 95, 3859–3867 (2000).

38. Apasov SG, Sitkovsky MV. The extracellular versus intracellular mechanisms of inhibition of TCR-triggered activation in thymocytes by adenosine under conditions of inhibited adenosine deaminase. Int Immunol 11, 179–189 (1999).

39. Cekic C, Sag D, Day YJ, Linden J. Extracellular adenosine regulates naive T cell development and peripheral maintenance. J Exp Med 210, 2693–2706 (2013).

40. Schiedel AC, Lacher SK, Linnemann C, Knolle PA, Muller CE. Antiproliferative effects of selective adenosine receptor agonists and antagonists on human lymphocytes: evidence for receptor-independent mechanisms. Purinergic Signal 9, 351–365 (2013).

41. Mastelic-Gavillet B, et al. Adenosine mediates functional and metabolic suppression of peripheral and tumor-infiltrating CD8(+) T cells. J Immunother Cancer 7, 257 (2019).

42. Mager LF, et al. Microbiome-derived inosine modulates response to checkpoint inhibitor immunotherapy. Science 369, 1481–1489 (2020).

43. Klysz DD, et al. Inosine induces stemness features in CAR-T cells and enhances potency. Cancer Cell 42, 266–282 e268 (2024).

44. Niemann B, et al. Apoptotic brown adipocytes enhance energy expenditure via extracellular inosine. Nature 609, 361–368 (2022).

45. Xu X, Chen Z, Bartman CR, Xing X, Olszewski K, Rabinowitz JD. One-carbon unit supplementation fuels purine synthesis in tumor-infiltrating T cells and augments checkpoint blockade. Cell Chem Biol 31, 932–943 e938 (2024).

46. Waldman AD, Fritz JM, Lenardo MJ. A guide to cancer immunotherapy: from T cell basic science to clinical practice. Nat Rev Immunol 20, 651–668 (2020).

47. Ohta A, et al. A2A adenosine receptor protects tumors from antitumor T cells. Proc Natl Acad Sci U S A 103, 13132–13137 (2006).

48. Cekic C, Linden J. Purinergic regulation of the immune system. Nat Rev Immunol 16, 177–192 (2016).

49. Chang CH, et al. Posttranscriptional control of T cell effector function by aerobic glycolysis. Cell 153, 1239–1251 (2013).

50. Ma S, Ming Y, Wu J, Cui G. Cellular metabolism regulates the differentiation and function of T-cell subsets. Cell Mol Immunol 21, 419–435 (2024).

51. Boison D. Adenosine kinase: exploitation for therapeutic gain. Pharmacological Reviews 65, 906–943 (2013).

52. Li L, Liu J, Delohery T, Zhang D, Arendt C, Jones C. The effects of teriflunomide on lymphocyte subpopulations in human peripheral blood mononuclear cells in vitro. J Neuroimmunol 265, 82–90 (2013).

53. Scherer S, et al. Pyrimidine de novo synthesis inhibition selectively blocks effector but not memory T cell development. Nat Immunol 24, 501–515 (2023).

54. Wang N, et al. AMPK-a key factor in crosstalk between tumor cell energy metabolism and immune microenvironment? Cell Death Discov 10, 237 (2024).

55. Hoxhaj G, et al. The mTORC1 Signaling Network Senses Changes in Cellular Purine Nucleotide Levels. Cell Rep 21, 1331–1346 (2017).

56. Mentch SJ, et al. Histone Methylation Dynamics and Gene Regulation Occur through the Sensing of One-Carbon Metabolism. Cell Metab 22, 861–873 (2015).

57. Lucas-Hourani M, et al. Inhibition of pyrimidine biosynthesis pathway suppresses viral growth through innate immunity. PLoS Pathog 9, e1003678 (2013).

58. Wiebking V, et al. Metabolic engineering generates a transgene-free safety switch for cell therapy. Nat Biotechnol 38, 1441–1450 (2020).

59. Yamamoto T, Koyama H, Kurajoh M, Shoji T, Tsutsumi Z, Moriwaki Y. Biochemistry of uridine in plasma. Clin Chim Acta 412, 1712–1724 (2011).

60. Battaglia S, et al. Uridine and pyruvate protect T cells’ proliferative capacity from mitochondrial toxic antibiotics: a clinical pilot study. Sci Rep 11, 12841 (2021).

61. Thompson JT, Wood DM, Dargan PI. Review of the fluoropyrimidine antidote uridine triacetate. Br J Clin Pharmacol 91, 615–627 (2025).

62. Hau RK, Wright SH, Cherrington NJ. Addressing the Clinical Importance of Equilibrative Nucleoside Transporters in Drug Discovery and Development. Clin Pharmacol Ther 114, 780–794 (2023).

63. Huang W, Sundquist K, Sundquist J, Ji J. Use of dipyridamole is associated with lower risk of lymphoid neoplasms: a propensity score-matched cohort study. Br J Haematol 196, 690–699 (2022).

64. Takeda A, et al. Single-Cell Survey of Human Lymphatics Unveils Marked Endothelial Cell Heterogeneity and Mechanisms of Homing for Neutrophils. Immunity 51, 561–572 e565 (2019).

65. Eberhardt J, Santos-Martins D, Tillack AF, Forli S. AutoDock Vina 1.2.0: New Docking Methods, Expanded Force Field, and Python Bindings. J Chem Inf Model 61, 3891–3898 (2021).

66. Maier JA, Martinez C, Kasavajhala K, Wickstrom L, Hauser KE, Simmerling C. ff14SB: Improving the Accuracy of Protein Side Chain and Backbone Parameters from ff99SB. J Chem Theory Comput 11, 3696–3713 (2015).

67. Wang J, Wolf RM, Caldwell JW, Kollman PA, Case DA. Development and testing of a general amber force field. J Comput Chem 25, 1157–1174 (2004).

68. Jakalian A, Jack DB, Bayly CI. Fast, efficient generation of high-quality atomic charges. AM1-BCC model: II. Parameterization and validation. J Comput Chem 23, 1623–1641 (2002).

69. Abraham MJ, et al. GROMACS: High performance molecular simulations through multi-level parallelism from laptops to supercomputers. SoftwareX 1**-****2**, 19–25 (2015).

70. Valdes-Tresanco MS, Valdes-Tresanco ME, Valiente PA, Moreno E. gmx_MMPBSA: A New Tool to Perform End-State Free Energy Calculations with GROMACS. J Chem Theory Comput 17, 6281–6291 (2021).

