## Supplemental Tables and Figures for "Adenosine impairs T cell function via activation of purine salvage pathway and AMP-induced inhibition of pyrimidine nucleotide biosynthesis"

Supplementary data for the manuscript: **Adenosine impairs T cell function via activation of purine salvage pathway and AMP-induced inhibition of pyrimidine nucleotide biosynthesis**

**Karolina Losenkova[1], Ursula J. Von Danwitz[1,2], Nora Kreisig[1,2], Samuel Svärd[1], Julia S. Baier[1,2], Corinna Weiler[1,2], Anton Zavialov[1], Neil J. Thomson[3], Christa E. Müller[2], Osman Gani[3], Sirpa Jalkanen[1], Akira Takeda[1*], Gennady G. Yegutkin[1*]**

[1]MediCity Research Laboratory and InFLAMES Flagship, University of Turku, Turku, Finland;
[2]PharmaCenter Bonn, Pharmaceutical Institute, Pharmaceutical & Medicinal Chemistry University of Bonn, Bonn, Germany
[3]Department of Pharmacy, Section for Pharmaceutical Chemistry, University of Oslo, Oslo, Norway.

**Supplementary Table 1.** Substrate selectivity of the OMPD active site

| Compound | MW | Charge | #$PO_4$ | Score | Assessment |
|---|---|---|---|---|---|
| UMP | 322 | −2 | 1 | −6.6 | Crystal pose validated; product of OMPD reaction |
| OMP | 365 | −3 | 1 | −7.1 | Crystal pose validated; natural substrate of OMPD |
| AMP | 345 | −2 | 1 | −7.7 | Modeled pose validated; inhibitor under investigation |
| ATP | 503 | −4 | 3 | −7.4 | Excluded: triphosphate tail incompatible with pocket; excess charge |
| ADP | 424 | −3 | 2 | −7.6 | Excluded: diphosphate tail too bulky; charge not compensated |
| ADO | 267 | 0 | 0 | −7.0 | Excluded: lacks phosphate anchor for Lys281/Lys314 salt bridges |
| GTP | 519 | −4 | 3 | −6.9 | Excluded: triphosphate tail incompatible with pocket; excess charge |
| GMP | 361 | −2 | 1 | −7.3 | Not excluded by docking; correct size/charge but no experimental evidence of binding or inhibition of OMPD |
| CMP | 321 | −2 | 1 | −7.4 | Not excluded by docking alone; correct size/charge |
| IMP | 346 | −2 | 1 | −7.1 | Not excluded by docking alone; correct size/charge |

Docking screen of different nucleotides and ADO into the OMPD active site (PDB 2QCD, chain A). Scores in kcal/mol (AutoDock Vina). Molecular weight (MW) is given for the ionic species at the stated formal charge.  Green: included in MD/FEP study. Red: excluded on structural or physicochemical grounds. Yellow: excluded based on other published and experimental data (see main text for details); and also on steric and electrostatic grounds despite favorable docking scores, since Vina scoring does not resolve the charge and tail-placement constraints of this catalytic pocket.

**Supplementary Table 2.** Primary antibodies used for fluorescence microscopy and flow cytometry

| Target | Antibody (clone, catalogue number) | Host | Manufacturer | Application |
|---|---|---|---|---|
| CD39 | Anti-CD39 Ab (hN1-$1_C$) | Guinea pig | Ectonucleotidases[1] | IF |
| CD73 | Anti-CD73 Ab (h5NT-$1_L$) | Rabbit | Ectonucleotidases[1] | IF |
| CD39 | BV711™ anti-CD39 Ab (clone A1, Cat #328228) | Mouse | Biolegend® | FACS |
| CD73 | PE-CF593 anti-CD73 Ab (clone AD2, Cat #562817) | Mouse | BD Horizon™ | FACS |
| ENT1 | Alexa Fluor® 647 anti-ENT1 Ab (ab311105) | Rabbit | Abcam | IF, FACS |
| CD3 | Alexa Fluor® 488 anti-CD3 Ab (clone HIT3a, Cat #300320) | Mouse | Biolegend® | IF, FACS |
| CD4 | PE anti-CD4 Ab (clone RPA-T4, Cat #555347) | Mouse | BD Pharmingen™ | FACS |
| CD8a | BV785™ anti-CD8a Ab (clone RPA-T8, Cat #301045) | Mouse | Biolegend® | FACS |
| PD-1 | BV650™ anti-PD-1 Ab (clone EH12.2H7, Cat #329950) | Mouse | Biolegend® | FACS |
| CD25 | PerCP-Cy5.5 anti-CD25 Ab (clone BC96, Cat #3026269) | Mouse | Biolegend® | FACS |
| GranzB | PE-Cy7 anti-GranzB Ab (clone QA16A02, Cat #372213) | Mouse | Biolegend® | FACS |
| IFN-γ | BV785™ anti-IFN-γ Ab (clone 4S.B3, Cat #502541) | Mouse | Biolegend® | FACS |
| Vimentin | Anti-vimentin Ab (919101) | Chicken | Biolegend® | IF |

[1] http://ectonucleotidases-ab.com/ Abbreviations: Ab, antibody, BV, brilliant violet, ENT1, equilibrative nucleoside transporter-1, FACS, fluorescence-activated cell sorter, GranzB, Granzyme B, IF, immunofluorescence microscopy, IFN-γ, interferon-gamma, PD-1, programmed cell death protein-1.

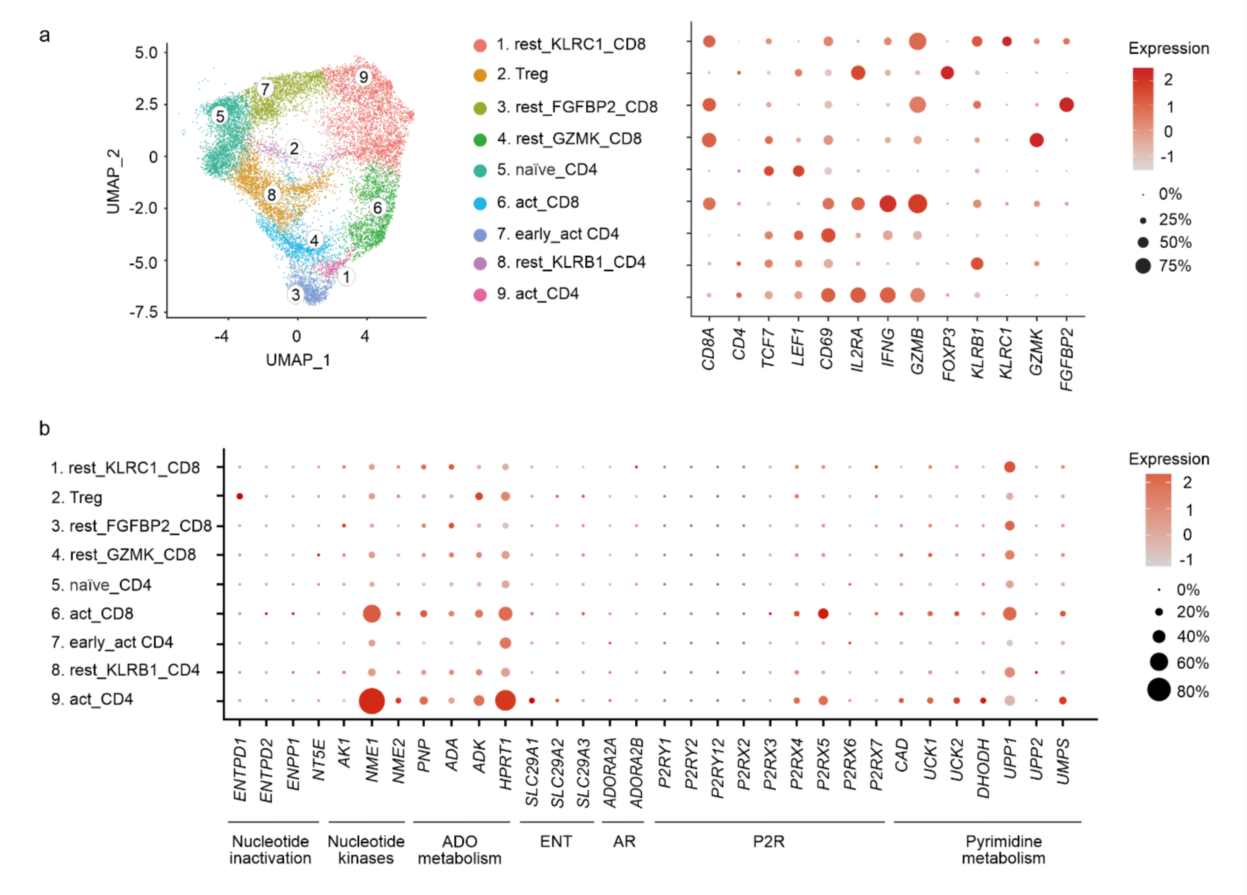


**Supplementary Fig. 1. scRNA-seq analysis of purine and pyrimidine metabolism in human T cells activated with anti-CD3/CD28 beads.** We utilized publicly available single-cell transcriptomic data of human T cells, which were isolated from the blood of two healthy volunteers and activated by incubating for 16 hours with human T-Activator CD3/CD28 beads [26]. (a) Unbiased clustering of these data identified 9 distinct clusters, distinguished by expression of CD4, CD8A, and markers of naïve (*TCF7, LEF1*), resting (*KLRC1, GZMK, FGFBP2*), activated (*CD69, IFNG, ILR2RA, GZMB*), and regulatory (*FOXP3*) T cells. (b) Compared with naïve and resting T cells, activated $CD8^+$ (cluster 6) and $CD4^+$ (cluster 9) T cells expressed higher levels of genes encoding purine- and pyrimidine-converting enzymes NDPK (especially the NDPK-A/*NME1* isoform), ADK (*ADK*), PNP (*PNP*), HPRT1, UCK1/2, and UMPS, as well as ENT1 (*SLC29A1*) and P2X4 (*P2RX4*) and P2X5 (*P2RX5*) receptors. The relative expression of the indicated genes is shown on a pseudocolor scale (log2(FC)), with the size of the dot representing the percentage of cells in T cell subset where the gene is detected.

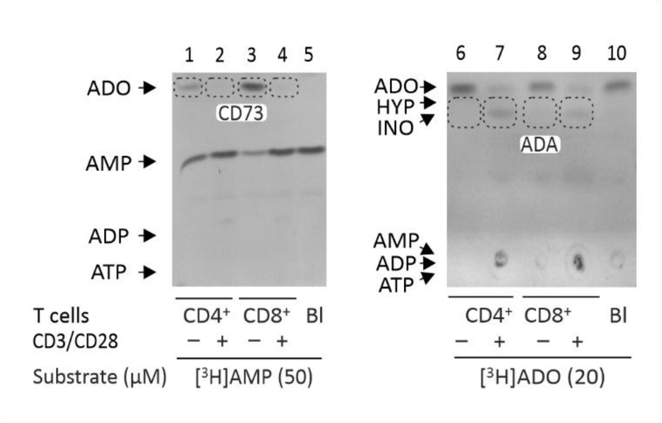


**Supplementary Fig. 2. Autoradiographic analysis of key ADO-metabolizing ectoenzymes in resting and activated CD4$^+$ and CD8$^+$ T cells.** CD3$^+$ T cells were isolated from the blood of healthy volunteers, separated into CD4$^+$ and CD8$^+$ T cell subsets by immunomagnetic selection, and incubated for four days without and with anti-CD3/CD28 beads, as indicated. Resting and activated T cells were incubated with 50 µM [$^3$H]AMP and 20 µM [$^3$H]ADO as respective substrates for CD73 and ADA. $^3$H-labeled substrates and their metabolites were separated by using two different solvent systems employed for the measurement of ecto-nucleotidase and ADA activities. Representative autoradiographic images of the patterns of [$^3$H]AMP (left) and [$^3$H]ADO (right) metabolism by CD4$^+$ and CD8$^+$ T cells are shown. The corresponding products of CD73 and ADA reactions, [$^3$H]ADO and [$^3$H]INO, are outlined by dashed boxes. The blanks (Bl) show the radiochemical purities of $^3$H-labelled substrates in the absence of T cells. Arrows indicate the positions of nucleotide standards, ADO, inosine (INO), and hypoxanthine (HYP).

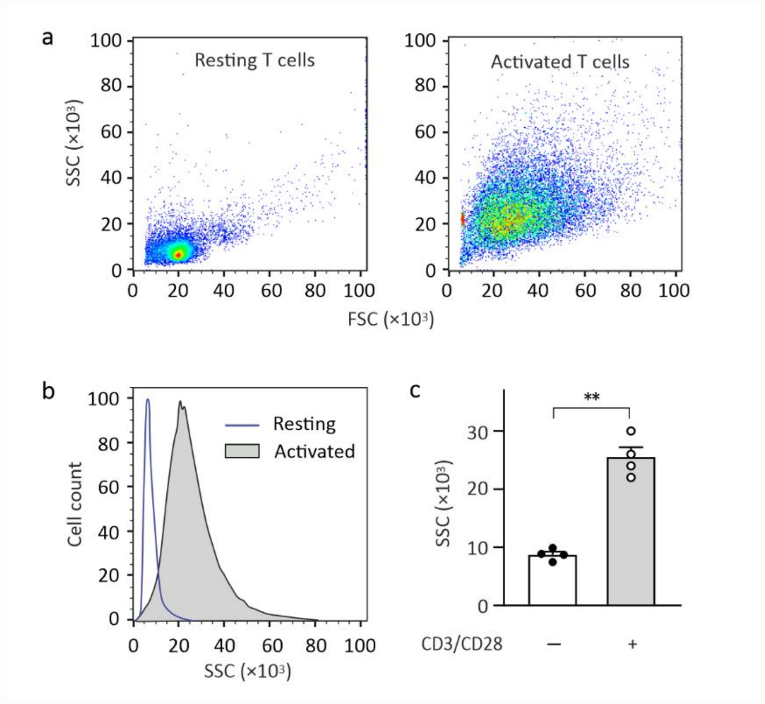


**Supplementary Fig. 3. Flow cytometric analysis reveals an increase in size and granularity of human T cells upon activation.** Human T cells were incubated for four days in the absence and presence of anti-CD3/CD28 beads. (a) The size and granularity of T cells were assessed using flow cytometry. Suspensions of the resting and activated T cells were displayed in forward scatter (FSC) versus side scatter (SSC) dot plots in logarithmic scale. (b) Representative normalized histograms of resting and activated T cells expressed as Cell Count versus SSC. (c) The graph shows the number of SSC events collected in the unstained resting and activated T cells from four independent experiments (mean + SEM). **P<0.01, determined by Student's t test (paired, two-tailed).

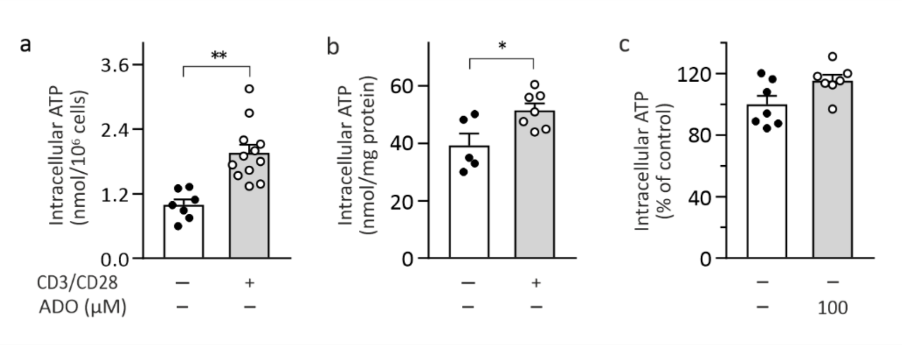


**Supplementary Fig. 4. Comparative analysis of intracellular ATP levels in resting and activated T cells.** T cells were isolated from the blood of healthy volunteers and incubated for four days without (-) and with (+) anti-CD3/CD28 beads, as indicated. (a,b) Resting and activated T cells were lysed and analyzed for intracellular ATP content using a bioluminescent luciferin-luciferase assay. Protein concentration in cell lysates was determined using the BCA Protein Assay Kit. The results are expressed as nanomoles of ATP per $10^6$ cells (a) or per milligram of cellular protein (b). Statistical significances were determined by unpaired Student's t-test (*P<0.05 and **P<0.01). (c) Bioluminescent analysis of intracellular ATP in resting T cells treated with 100 μM ADO. The results are plotted as the percentage of ATP levels measured in control vehicle-treated cells.

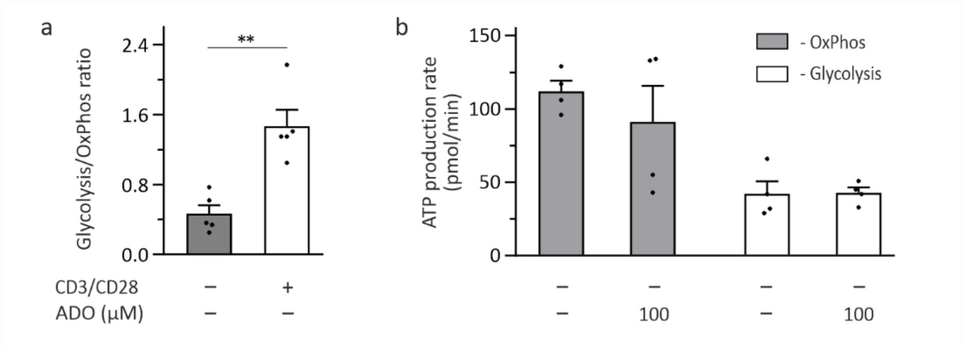


**Supplementary Fig. 5. ADO treatment does not affect conventional bioenergetic pathways in resting T cells.** T cells were isolated from the blood of healthy volunteers and incubated for four days without (-) and with (+) anti-CD3/CD28 beads, as indicated. T cell bioenergetics was determined using the Agilent Seahorse XF Cell Mito Stress Kit. (a) Simultaneous measurements of the rates of oxygen consumption and extracellular acidification allowed to determine cellular ATP production through oxidative phosphorylation (OxPhos) and glycolysis. The results are expressed as glycolysis to OxPhos ratio from five independent experiments (mean + SEM). **P<0.01, determined by Student's t-test (paired, two-tailed). (b) Resting T cells were incubated for 24 hours with ADO (0-100 µM). ATP production rates through mitochondrial OxPhos and glycolysis were determined and expressed as picomoles of ATP per minute (mean + SEM, n=4).

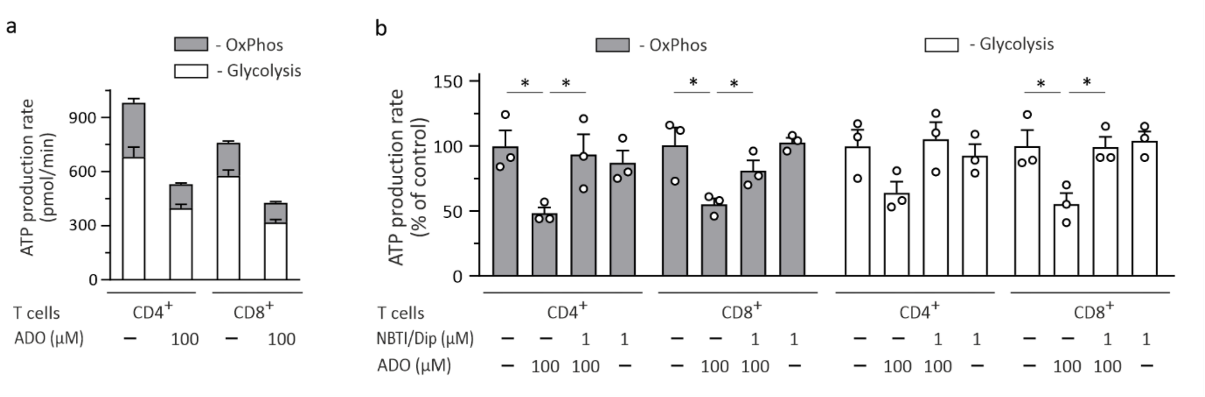


**Supplementary Fig. 6. ADO inhibits mitochondrial and glycolytic ATP production in activated CD4+ and CD8+ T cells via ENT1-dependent mechanism.** T cells were isolated from the blood of healthy volunteers, separated into CD4+ and CD8+ T cell subsets by immunomagnetic selection, and stimulated for four days with anti-CD3/CD28 beads. Activated CD4+ and CD8+ T cells were treated for 24 hours with the indicated concentrations of ADO and ENT1 inhibitors NBTI and dipyridamole (NBTI/Dip). T cell bioenergetics was evaluated using the Agilent Seahorse XF Cell Mito Stress Kit. (a) ATP production rates in CD4+ and CD8+ T cells under basal conditions and after addition of 100 μM ADO. Total ATP production was calculated as the sum of the ATP production rates from glycolysis and oxidative phosphorylation (OxPhos). The results from one representative experiment performed with four technical replicates are shown. (b) ADO inhibits ATP production rates in the activated CD4+ and CD8+ T cells, which can be rescued by blocking cellular nucleoside uptake with NBTI and dipyridamole. The results are presented as mean + SEM. *P<0.05, determined by Student's t test (unpaired, two-tailed).

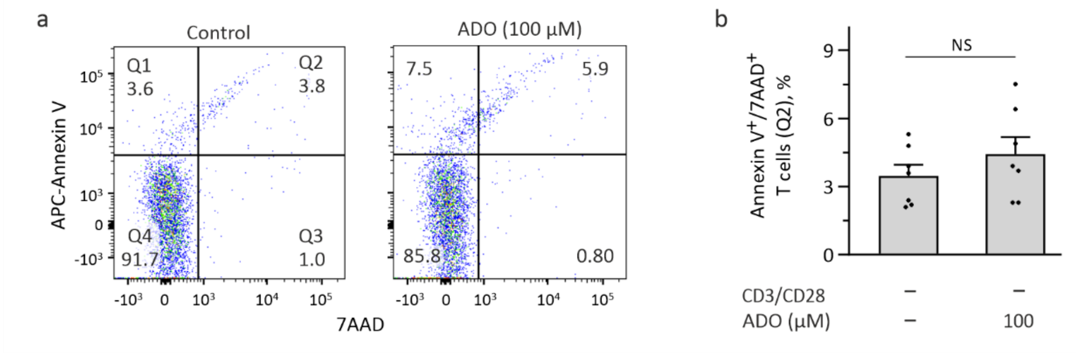


**Supplementary Fig. 7. ADO treatment does not affect the viability of human naïve T cells.** T cells were isolated from the blood of healthy volunteers, maintained in a resting state by culturing for four days in T cell medium containing IL-7, and incubated for additional 24 hours without and with 100 μM ADO. The viability of T cells was assessed using the APC Annexin V Apoptosis Detection Kit with 7AAD. (a) Representative dot plots for Annexin V versus 7AAD staining of control and ADO-treated T cells. (b) The frequencies of annexin $V^+$/$7AAD^+$ apoptotic T cells (Q2) were quantified using FlowJo software and expressed as percentage of total T cell numbers (mean + SEM). No statistically significant differences in apoptosis rate were detected between the control and ADO-treated T cells. NS, non-significant.

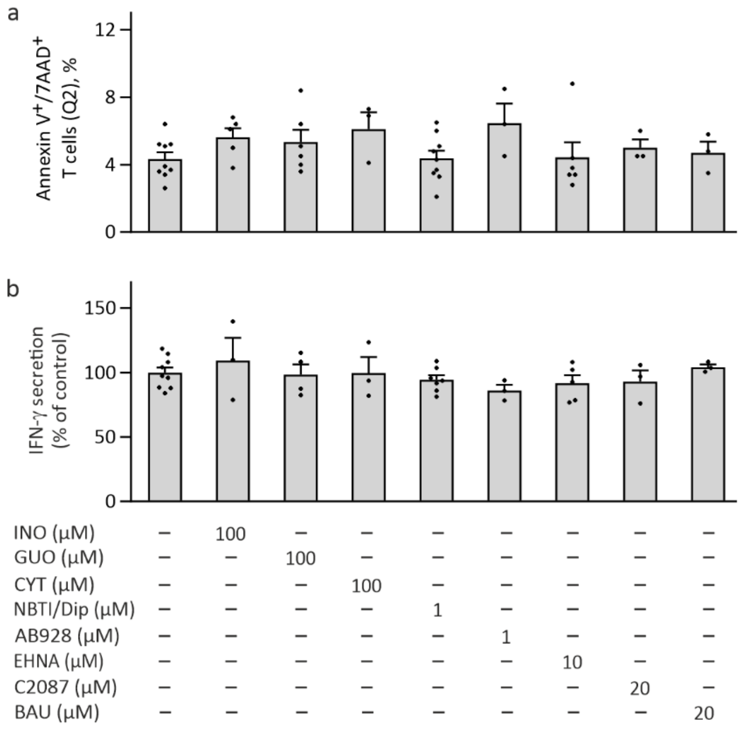


**Supplementary Fig. 8. The effect of nucleosides and other purinergic and pyrimidinergic drugs on T cell viability and IFN-γ production by activated T cells.** T cells were stimulated for four days with anti-CD3/CD28 beads, incubated for additional 24 hours with the indicated concentrations of purine and pyrimidine nucleosides and other compounds, and assessed for cell viability and effector function by flow cytometry. The frequencies of annexin $V^+$/$7AAD^+$ apoptotic T cells (a) and mean fluorescence intensities of IFN-$\gamma^+$ T cells (b) were quantified using FlowJo (mean + SEM). Statistical significances were determined by one-way ANOVA with Dunnett's multiple comparison test, with no statistical differences being detected between the control and treated T cells.

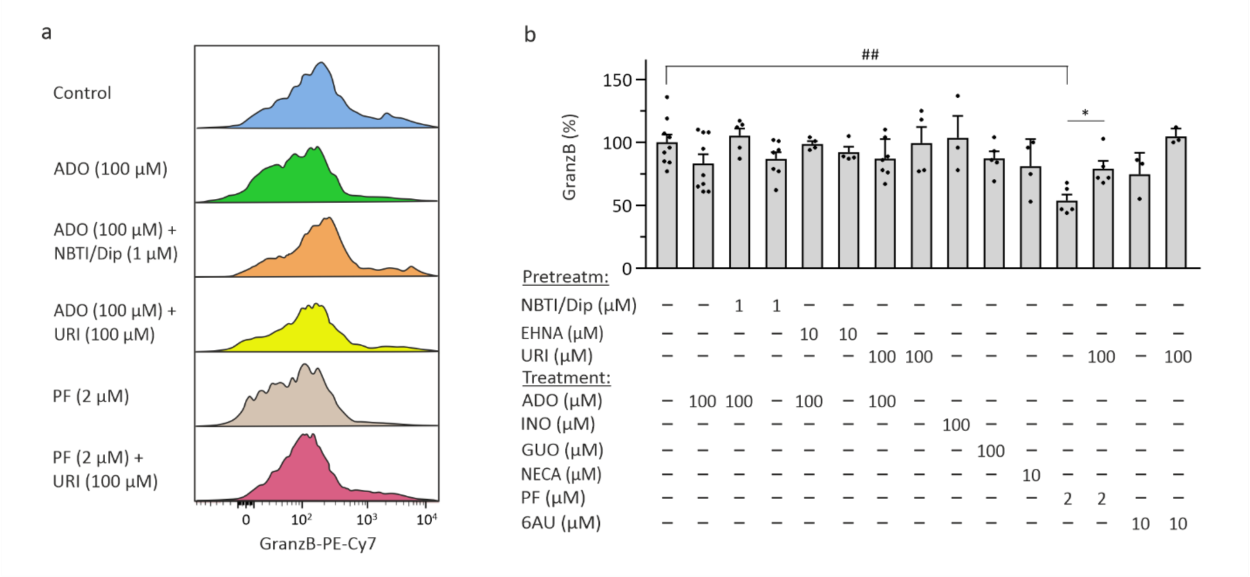


**Supplementary Fig. 9. The effect of ADO and other drugs on the production of Granzyme B by activated T cells.** T cells were stimulated for four days with anti-CD3/CD28 beads and incubated for additional 24 hours with 100 µM ADO and other compounds, as outlined in Figure 3a. Treated T cells were subsequently stained with PE-Cy7-conjugated anti-Granzyme B (GranzB) antibody, and analyzed by flow cytometry. (a) The representative histograms show the expression levels of GranzB in control and treated T cells. (b) Mean fluorescence intensities of $GranzB^+$ T cells were quantified using FlowJo and expressed as percentage of control (mean + SEM). Statistical significances were determined by one-way ANOVA with Dunnett's multiple comparison test ($^{\#\#}P<0.01$), or unpaired Student's t-test (*P<0.05).

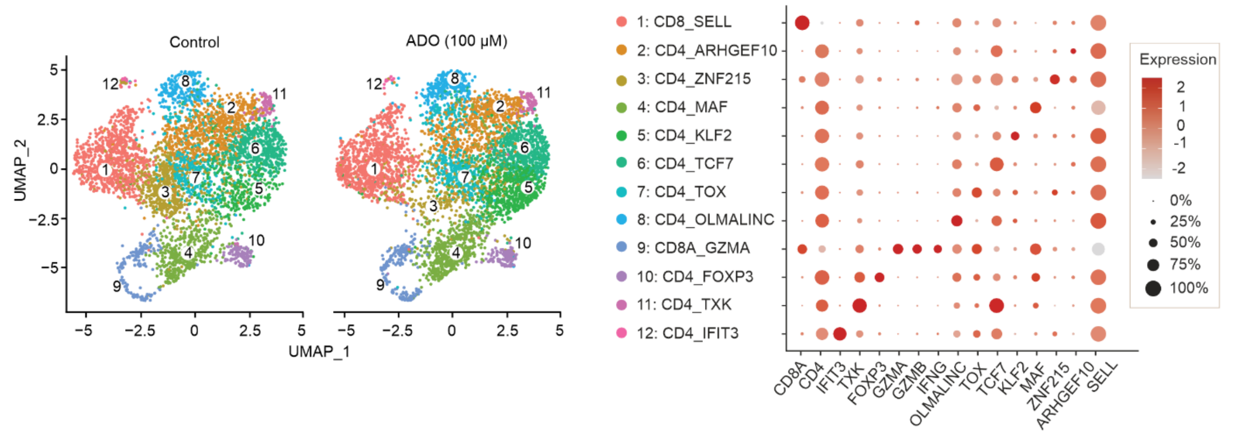


**Supplementary Fig. 10. scRNA-seq analysis of activated human T cells treated with ADO.** T cells from two donors were activated for four days with anti-CD3/CD28 beads and incubated for additional 24 hours in the absence (Control) or presence of 100 uM ADO. After treatment, T cells were separately processed for scRNA-seq and combined for downstream analysis. Transcriptome-based clustering of the single-cell expression profiles revealed 12 clusters, with no significant changes in cluster composition observed between the control and ADO-treated cells. CD8 was detected in two clusters, which were identified as naïve $CD8^+$ T cells, hallmarked by high expression of L-selectin (SELL) (cluster 1), and $CD8^+$ effector T cells, characterized by high expression of IFN-γ (IFNG) and Granzymes A (GZMA) and B (GZMB) (cluster 9). Among the $CD4^+$ T cell subsets, cluster 4 displayed high expression of MAF basic-leucine zipper transcription factor (MAF), which is a characteristic transcription factor of effector $CD4^+$ T cells. Cluster 10 expressed FOXP3, identifying these cells as Treg cells. Dot plots show the expression of representative genes differentially expressed among T cell clusters. The relative expression of the indicated genes is shown on a pseudocolor scale (log2(FC)), with the size of the dot representing the percentage of cells in T cell subset where the gene is detected. Other abbreviations: *ARHGEF10,* Rho guanine nucleotide exchange factor 10; *IFIT3*, Interferon-induced protein with tetratricopeptide repeats 3; *OLMALINC*, oligodendrocyte maturation-associated long intergenic non-coding RNA; *TOX*, thymocyte selection-associated HMG BOX; *TXK,* tyrosine-protein kinase TXK; *ZNF215*, Zinc finger protein 215.

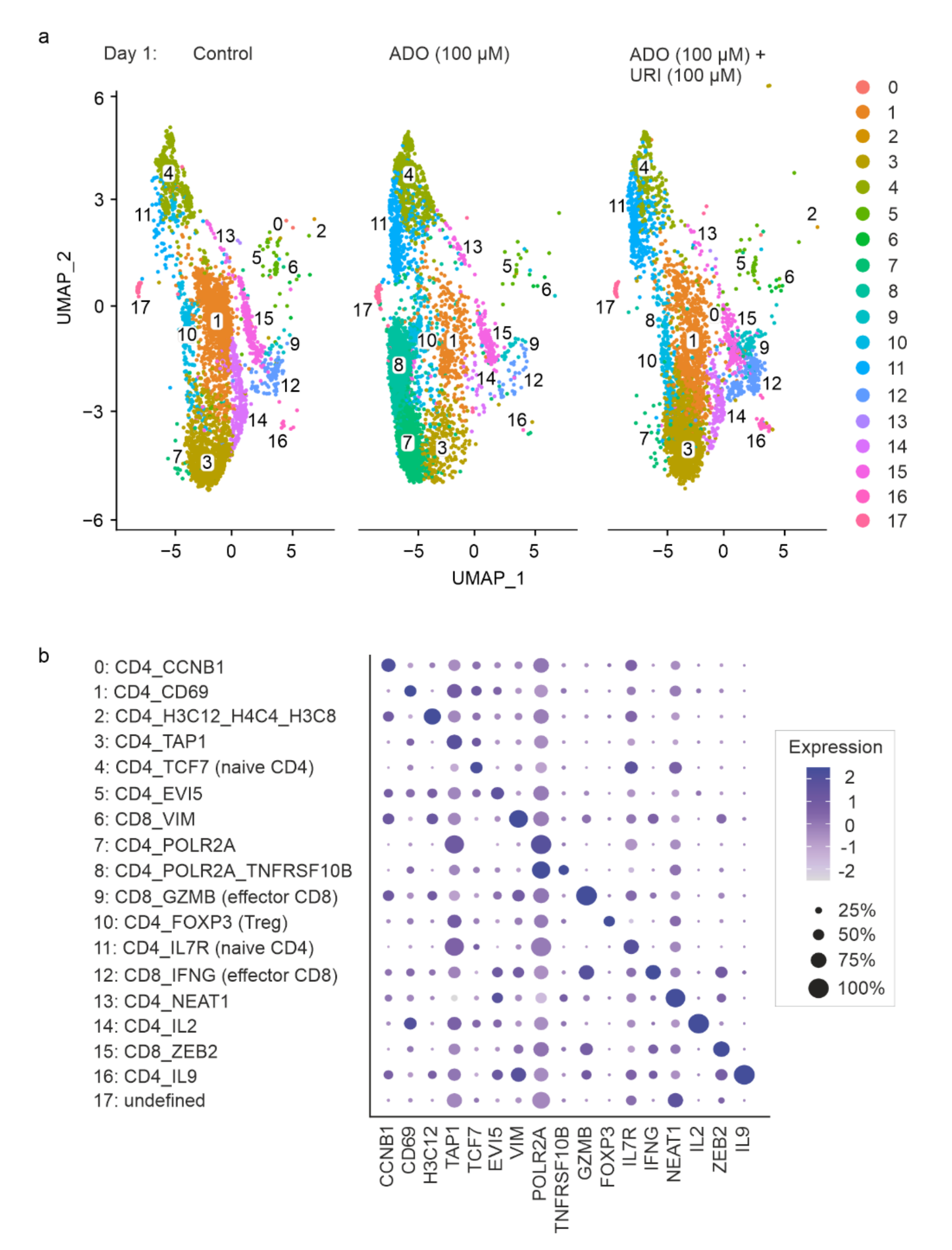


**Supplementary Fig. 11. scRNA-seq analysis of human T cells activated with anti-CD3/CD28 beads in the presence of ADO and uridine.** Peripheral blood T cells from two healthy adult volunteers were incubated with anti-CD3/CD28 beads in the absence (control) and presence of 100 µM ADO and uridine (URI). The cells were treated for one (panel a) or three (Fig. 6a) days, processed for scRNA-seq, and combined for downstream analysis. (b) Transcriptome-based clustering of the single-cell expression profiles revealed 17 clusters. Dot plots show the expression of representative genes differentially expressed among T cell clusters, including genes encoding G2/mitotic-specific cyclin-B1 (*CCNB1*), early activation marker CD69, replication-dependent histone H3 protein (*H3C12*), ATP Binding Cassette Subfamily B Member Transporter 1 (*TAP1*), DNA-directed RNA polymerase II subunit RPB1 (*POLR2A*), and TNF receptor superfamily member 10B (*TNFRSF10B*). The relative expression of the indicated genes is shown on a pseudocolor scale (log2(FC)), with the size of the dot representing the percentage of cells in T cell subset where the gene is detected. Other abbreviations: *EVI5*, Ecotropic viral integration site 5; *FOXP3,* Forkhead box P3; *NEAT1*, Nuclear enriched abundant transcript 1; *TCF7*, Transcription factor 7; *VIM*, vimentin; *ZEB2*, Zinc finger E-box binding protein 2.

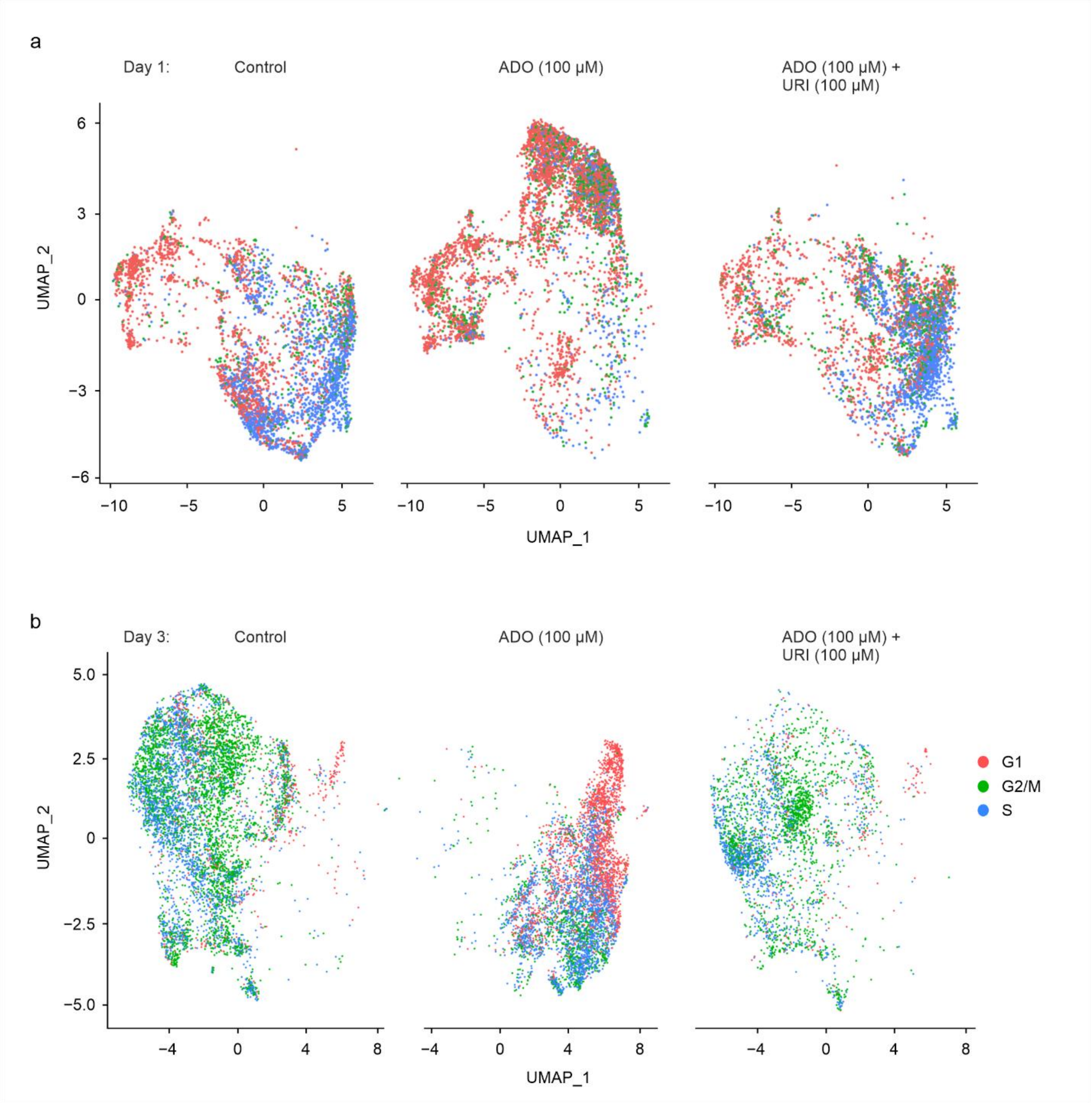


**Supplementary Fig. 12. UMAP visualization of human T cells activated with anti-CD3/CD28 beads for one (a) and three (b) days in the presence of ADO and uridine.** Cell cycle phase in each cell was calculated based on its expression of G2/M and S phase markers.

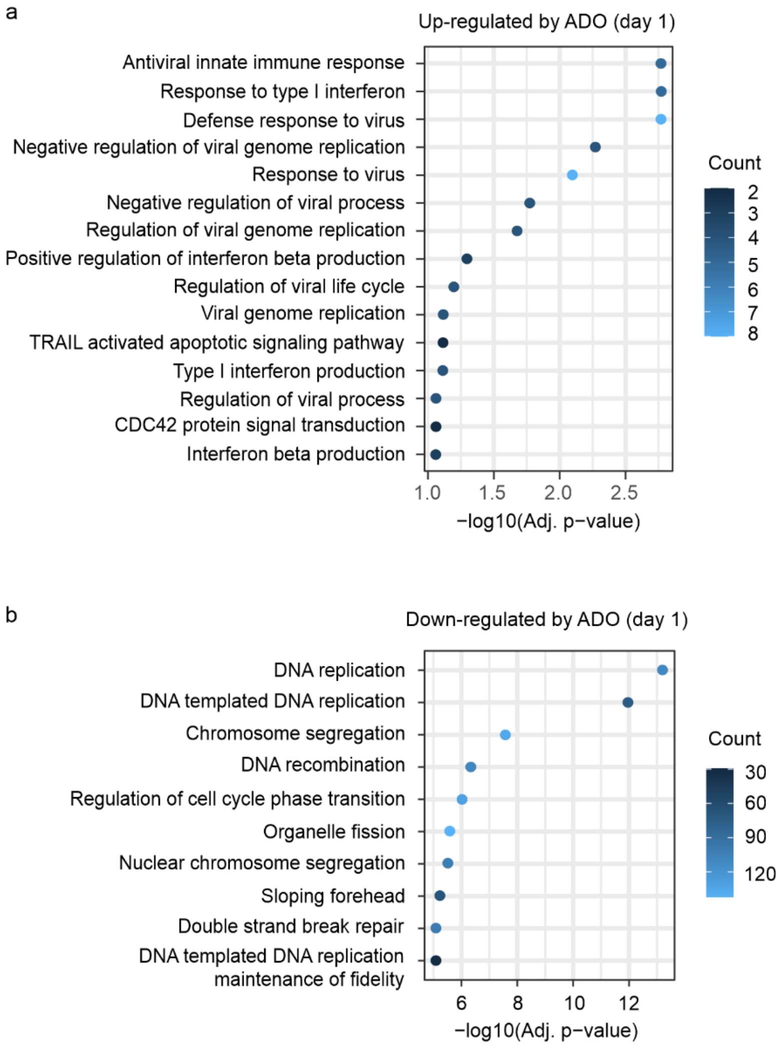


**Supplementary Fig. 13. scRNA-seq analysis of human T cells activated for one day with anti-CD3/CD28 beads in the presence of ADO.** Peripheral blood T cells were isolated from two healthy volunteers and incubated with anti-CD3/CD28 beads in the absence and presence of 100 µM ADO. After treatment, T cells were separately processed for scRNA-seq and combined for downstream analysis. Gene Ontology annotation enrichment analysis shows the top genes up-regulated (a) and down-regulated (b) in ADO-treated T cells after one day of activation, determined using the EnrichR package. The color of the dots represents the number of enriched genes. FDR-corrected p values are shown.
